# Cell-type-specific striatal modulation of amygdalar acetylcholine in salience assignment

**DOI:** 10.1101/2024.07.30.605919

**Authors:** Aixiao Chen, Yunjing Li, Hangfei Zhu, Hanmei Gu, Yanhong Weng, Qinyong Ye, Wuqiang Guan, Qingtao Sun, Bo Li, Fuqiang Xu, Hanfei Deng, Xiong Xiao

## Abstract

The salience assignment is pivotal for both natural and artificial intelligence. Pioneering studies established that basal forebrain cholinergic neurons process behaviorally relevant salient information. However, the neural circuit mechanism underlying salience assignment remains poorly understood. Here we show that the acetylcholine (ACh) level in the basolateral amygdala (BLA) dynamically represented behavioral salience. Distinct neuronal subpopulations in the nucleus accumbens (NAc), D1- and D2-expressing medium spiny neurons (MSNs), antagonistically and specifically promote and suppress ACh release in the BLA, but not the cortex and hippocampus. These striatal D1 and D2 MSNs regulate BLA ACh by disinhibiting and inhibiting cholinergic neurons in the basal forebrain subregion substantia innominata (SI), respectively. Optogenetic manipulations of the pathway from striatal D1 and D2 MSNs to the SI opposingly affect associative learning. Our findings uncover an unconventional role of striatal MSNs in salience assignment via regulating the salience-representing amygdalar ACh activity.

## INTRODUCTION

The brain receives a constant influx of complex sensory inputs from the environment. Salient stimuli, distinguished by their physical distinctiveness, relevance, or emotional impact, are easily noticed, perceived, and remembered. Behavioral salience, the property by which certain stimuli stand out, enables the brain to efficiently allocate its limited perceptual and cognitive resources to the most behaviorally relevant stimuli ^1,2^. In the ever-changing environment where our goals and internal states also fluctuate, the stimuli salience can change swiftly. Salience assignment mechanisms allow the brain to dynamically adjust attentional priorities in response to shifting environmental demands, internal states, and goals. This process optimizes the allocation of resources to the most pertinent information for the intended behavior, such as perception, attentional control, and decision-making ^2–4^. In the realm of artificial intelligence, attention mechanisms akin to salience assignment have propelled recent advances by enhancing the ability of transformer models to extract relevant signals and suppress noise within vast datasets ^5,6^. However, the neural mechanism underlying salience assignment in the brain remains elusive.

Previous studies have shown that acetylcholine (ACh) is a key neuromodulator critical for conscious awareness, salience, and attentional control, and is essential for associative learning and memory ^7–15,15,16,16–19^. Cholinergic deficiencies represent a prominent feature of attention-related impairments observed in Alzheimer’s disease and Parkinson’s dementia ^20,21^. Selective lesion of cholinergic neurons, pharmacological blockade of ACh receptors, or inhibition of cholinergic neural activity all result in impairment of sensory discrimination, attention, memory, and conditioning performance ^16,22–24^. Activation of cholinergic neurons has been shown to facilitate sensory discrimination and associative learning ^16,18,25^. Cholinergic neurons in the basal forebrain also respond to both positive and negative reinforcers ^11–13,26^.

The basal forebrain cholinergic neurons project predominantly to three main targets: the cerebral cortex, hippocampus, and basolateral amygdala (BLA) ^27–29^. Stimuli that evoke strong emotions, or are associated with values are more likely to be perceived as salient. The BLA is particularly pivotal in processing emotional and motivational information, associating cues with valenced outcomes, and maintaining emotional salience ^27,30–33^. The principal neurons in the BLA have been shown to encode both positive and negative values of stimuli ^32,34^. Recent studies have begun to reveal the essential role of BLA ACh in cue-reward learning, fear conditioning, and reinforcement prediction error ^12,13,15,25,35^. However, the coding of various forms of behaviorally relevant salience in the BLA ACh has not been systematically investigated. Importantly, a fundamental question is how the behavioral salience, which is represented by the ACh level, is regulated and contributes to associative learning.

To address how the brain solves the salience-assignment problem, we attempted to identify the neural circuit that could tailor the ACh level in the BLA. We first recorded dynamic changes of ACh release in the BLA during the mouse behavior. We demonstrated that BLA ACh is robustly activated by a variety of behaviorally salient events, including novel (“unfamiliar”) stimuli, reinforcers and their predicting cues that gain salience, dependent on animals’ learning experience, and changes in task engagement and homeostatic states. We further identified that different neuronal subpopulations in the striatal region nucleus accumbens (NAc), D1- and D2-receptor-expressing medium spiny neurons (MSNs), opposingly regulate the release of BLA ACh and affect associative learning, via disinhibiting and inhibiting the basal forebrain cholinergic neurons in the substantia innominata (SI), respectively. Our findings uncover a unique role for the basal ganglia in regulating the salience assignment, a process crucial for adaptive and efficient associative learning.

## RESULTS

### BLA ACh encodes multiple forms of behavioral salience

We set out to systematically characterize the *in vivo* response properties of BLA ACh to a battery of behaviorally salient events. For this purpose, we injected adeno-associated virus (AAV) expressing the genetically encoded fluorescent ACh sensor (GRAB-ACh3.8m, abbreviated as ACh3.8m) ^36^ into the BLA, and then used fiber photometry to record intrinsic dynamics of the ACh signal in head-fixed behaving mice (Figure 1A). Behavioral salience can be influenced by various factors, such as novelty, stimulus intensity, and behavioral relevance ^37,38^.

**Figure 1.**
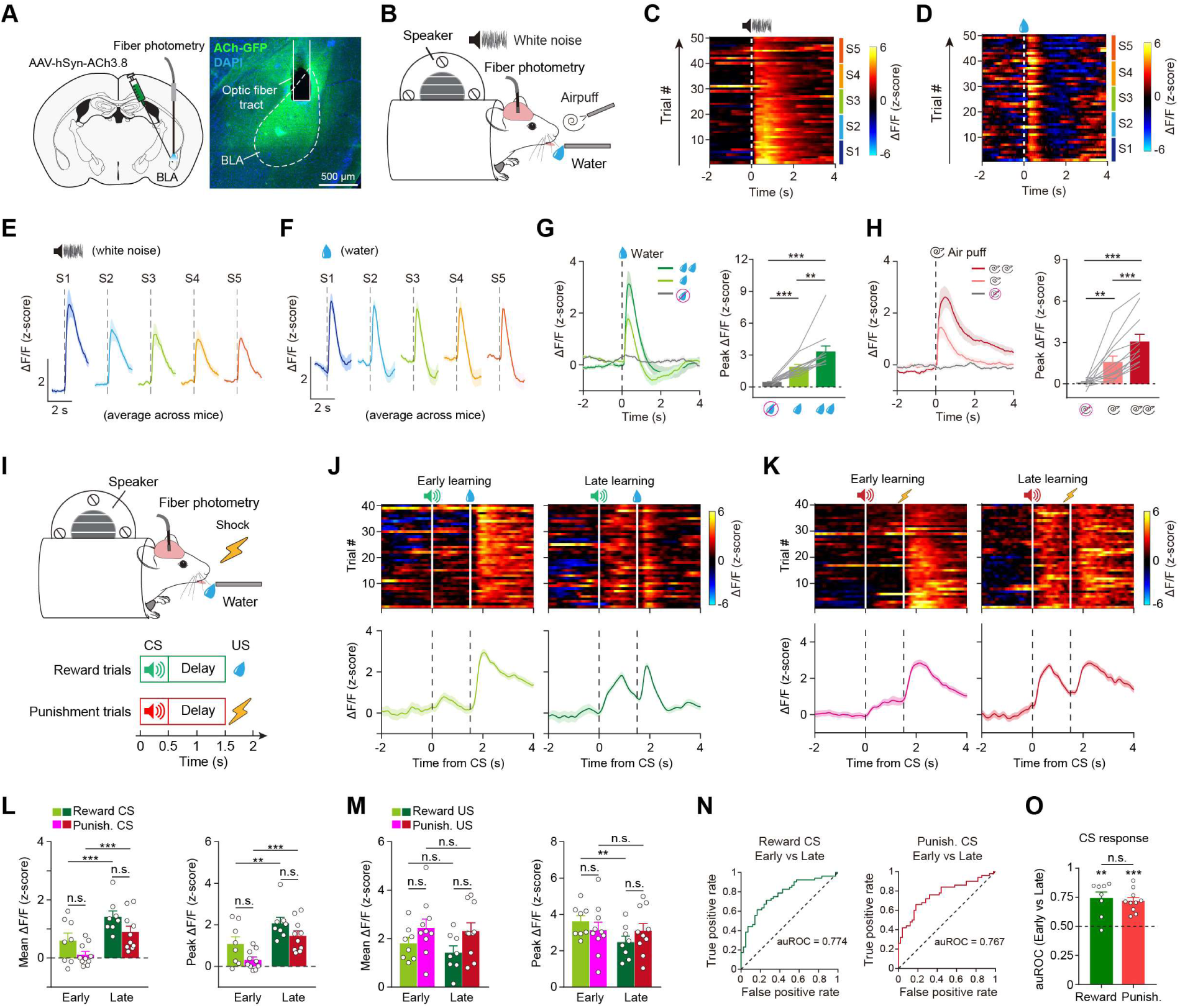
BLA ACh encodes behavioral salience irrespective of valence. **(A)** Left: schematic of the approach to record BLA ACh. Right: representative coronal image showing the expression of a fluorescent acetylcholine sensor (ACh3.8) in the BLA and the placement of optical fiber for photometry. **(B)** Schematics of the experimental setup. **(C)** Trial-by-trial heatmap of BLA ACh responses to the white noise in an example session. Each session consists of 50 trials, divided into 5 sections (10 trials per section). **(D)** Trial-by-trial heatmap of BLA ACh responses to the water in an example session. **(E)** Average BLA ACh responses (n = 12 mice) to white noise along sections. **(F)** Average BLA ACh responses (n = 12 mice) to water along sections. **(G)** Left: average BLA ACh responses (n = 12 mice) to different sizes of reward. Right: quantification of peak BLA ACh responses (F_(2, 33)_ = 18.93, p < 0.001, one-way ANOVA followed by Tukey’s test). **(H)** Left: average BLA ACh responses (n = 12 mice) to different sizes of punishment. Right: quantification of peak BLA ACh responses (F_(2, 33)_ = 14.4, p < 0.001, one-way ANOVA followed by Tukey’s test). **(I)** Schematics of the experimental setup (top) and the Pavlovian conditioning task design (bottom). **(J)** Trial-by-trial (top) and trial-averaged (bottom) BLA ACh responses in reward trials at the early (left) and late (right) stages of learning from a representative mouse in the Pavlovian task. **(K)** Trial-by-trial (top) and trial-averaged (bottom) BLA ACh responses in punishment trials at the early (left) and late (right) learning stages from a representative mouse in the Pavlovian task. **(L)** Quantification of the mean (left) (F_(1, 16)_ = 40.36, p < 0.001) and peak (right) (F_(1, 16)_ = 38.24, p < 0.001) BLA ACh responses to conditioned stimuli (CS) in the Pavlovian task. Two-way ANOVA followed by Bonferroni’s test. **(M)** Quantification of the mean (left) (F_(1, 16)_ = 3.07, p = 0.099) and peak (right) (F_(1, 16)_ = 7.42, p = 0.015) BLA ACh responses to unconditioned stimuli (US) in the Pavlovian task. Two-way ANOVA followed by Bonferroni’s test. **(N)** Receiver operating characteristic (ROC) curve evaluating the discriminability of BLA ACh responses to reward CS (left, auROC = 0.774) and punishment CS (right, auROC = 0.767) between early and late trials from a representative mouse. **(O)** Quantification of area under the ROC curve (auROC) for CS responses in reward and punishment trials (auROC vs 0.5, p = 0.002 for reward, p < 0.001 for punishment, paired t-test; reward vs punishment, p = 0.66, t-test). **p < 0.01, ***p < 0.001; n.s., non-significant (p > 0.05). Data are presented as mean ± SEM. Shaded areas represent SEM.

First, we presented water-restricted mice with white noise (novel stimulus) or water reward (Figure 1B). BLA ACh responses to the water delivery were robust and sustained across trials, whereas the responses evoked by white noise showed a gradual decline (Figures 1C-1F and S1A-S1D). Next, we presented mice with different magnitudes of reward (water) or punishment (air puff) (Figure S1E). BLA ACh responses scaled proportionally with the magnitude of the reward or punishment (Figures 1G, 1H, S1F, and S1H). Importantly, this ACh response profile did not reflect the rate of licking or eye blinking (Figures S1F-S1O), thus differed from the cortical ACh that tracks animals’ motor-categorized arousal levels ^39^.

To further test whether behavioral relevance affects BLA ACh responses, we trained mice to associate an auditory cue (conditioned stimulus, CS) with either an appetitive or aversive outcome (unconditioned stimulus, US) in a Pavlovian conditioning task (Figure 1I). The CS responses of BLA ACh increased significantly after learning (Figures 1J-1M). The CS responses can distinguish “early” versus “late” phases, but not as much for “reward” versus “punishment” cues (Figures 1N and 1O). Notably, cue responses of BLA ACh preceded the emergence of anticipatory licking during CS (Figures S2A-S2C) and did not track licking behavior (Figures 2D and 2E), resembling the temporal dynamics observed in dopamine signaling during associative learning ^40^.

**Figure 2.**
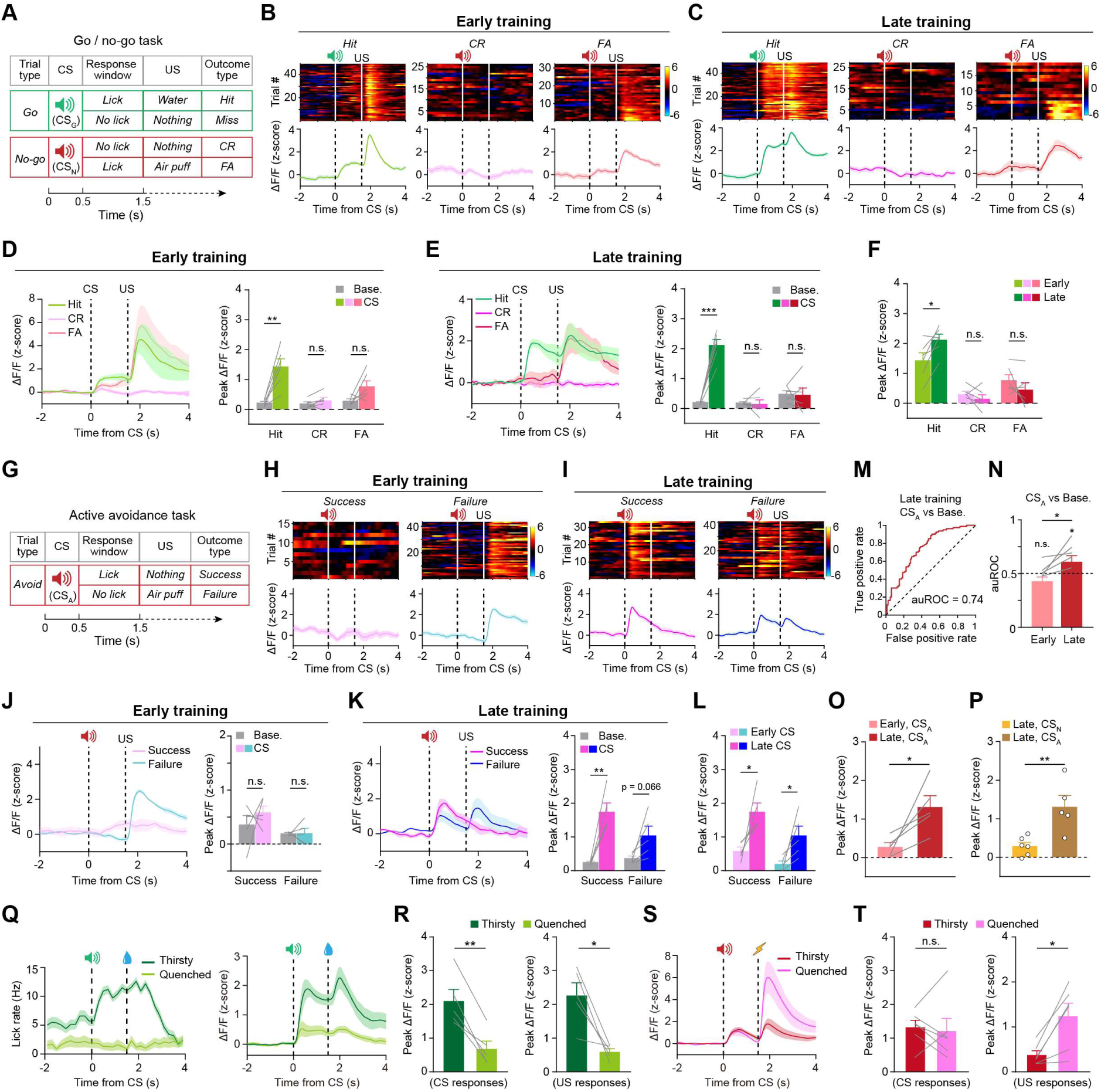
Modulation of salience response in BLA ACh by behavioral context and homeostatic state. **(A)** Schematic of the go/no-go task design. CR, correct rejection; FA, false alarm. CS_G_, CS for “go”; CS_N_, CS for “no-go”. **(B, C)** Trial-by-trial (top) and average (bottom) BLA ACh responses of a representative mouse at the early (B) and late (C) stages of the go/no-go task. **(D, E)** Average BLA ACh responses (left) and quantification of peak CS responses (right) for different trial outcomes at the early (D) (hit, p = 0.0039; CR, p = 0.22; FA, p = 0.071; paired t-test) and late (E) (hit, p < 0.001; CR, p = 0.75; FA, p = 0.87; paired t-test) stages of the go/no-go task. **(F)** Quantification of peak CS responses of BLA ACh in different trial outcomes at the early and late training stages of the go/no-go task (n = 5 mice; hit, p = 0.022; CR, p = 0.34; FA, p = 0.26; paired t-test). **(G)** Schematic of the active avoidance task design. CS_A_, CS for “avoid”. **(H, I)** Trial-by-trial (top) and average (bottom) BLA ACh responses of a representative mouse at the early (H) and late (I) stages of the active avoidance task. **(J, K)** Average BLA ACh responses (left) and quantification of peak CS responses (right) for different trial outcomes at the early (J) (success, p = 0.46; failure, p = 0.99; paired t-test) and late (K) (success, p = 0.004; failure, p = 0.066; paired t-test) stages of the active avoidance task. **(L)** Quantification of peak CS responses for different trial outcomes at the early and late training stages of the active avoidance task (n = 5 mice; success, p = 0.033; failure, p = 0.046; paired t-test). **(M)** ROC curve evaluating the discriminability of BLA ACh responses to CS_A_ and baseline at the late stage of the active avoidance task from a representative mouse. **(N)** Quantification of auROC for the early and late training stages of the active avoidance task (auROC vs 0.5, p = 0.41 for early, p = 0.015 for late; early vs late, p = 0.044; paired t-test). **(O)** Quantification of peak BLA ACh responses for CS_A_ at the early and late stages of the active avoidance task (p = 0.033, t-test). **(P)** Quantification of peak BLA ACh responses for CS_N_ and CS_A_ at the late stage of the go/no-go and the active avoidance task, respectively (p = 0.0058, t-test). **(Q)** Average lick rate (left) and BLA ACh responses (right) of a representative mouse during reward trials in the thirsty and quenched states in the Pavlovian task. **(R)** Quantification of the peak responses of BLA ACh for reward CS (left; p = 0.0037; paired t-test) and US (right; p = 0.0168; paired t-test) in the thirsty and quenched states (n = 5 mice). **(S)** Average BLA ACh responses of a representative mouse during punishment trials in the thirsty and quenched states in the Pavlovian task. **(T)** Quantification of the peak responses of BLA ACh for punishment CS (left; p = 0.7485; paired t-test) and US (right; p = 0.0136; paired t-test) in the thirsty and quenched states (n = 5 mice). *p < 0.05, **p < 0.01, ***p < 0.001; n.s., non-significant (p > 0.05). Data are presented as mean ± SEM. Shaded areas represent SEM.

Prediction error (PE), representing deviations from our expectation, is a salient signal that captures our attention and drives adaptive learning. To test whether BLA ACh can encode PE, we recorded BLA ACh in different settings of associative learning, pairing cues with probabilistic outcomes (Figure S2F). We found that the uncued (or surprised) reward evoked relatively larger ACh responses (Figures S2G, S2I, and S2J), which is a feature of PE. However, the surprised punishment did not induce larger responses of BLA ACh (Figures S2H, S2K, and S2L). These results indicate that BLA ACh is particularly sensitive to the salience of unexpected positive reinforcement events.

### BLA ACh encodes motivational salience in a context-dependent manner

The salience of behaviorally relevant stimuli can be shaped by animals’ motivation, influenced by both the external behavioral setting and their internal homeostatic state. To understand how behavioral context affects BLA ACh signals, we first trained mice to perform an operant “go/no-go” task, where mice need to lick to the “go” cue (CSG) to receive a reward and withhold licking to the “no-go” cue (CSN) to avoid punishment (Figure 2A). With learning, the ACh response to the CSG increased significantly, whereas the response to the CSN remained consistently low (Figures 2B-2F). Next, we trained mice to perform an active avoidance task, where mice need to lick in response to the “avoid” cue (CSA, the same auditory stimuli as the “no-go” cue) to avoid air puff delivery (Figure 2G). In the early learning period, the ACh response to the CSA was relatively weak (Figures 2H and 2J). Notably, the ACh response increased significantly to the CSA after learning (Figures 2I, 2K, and 2L-2O). In both tasks, we used the same auditory tone for “no-go” and “avoid” trials, yet we observed strikingly different ACh responses (Figure 2P). In the “no-go” trials, mice displayed passive “no-lick” behavior, whereas in the “avoid” trials, they exhibited heightened behavioral engagement. These observations strongly suggest that BLA ACh encoding of salience is contingent upon the motivational engagement of the animal.

To understand how homeostatic state affects BLA ACh signals, we examined the impact of thirsty versus quenched states on the CS- and US-evoked ACh activity. The cue predicting the availability of water elicited robust anticipatory licking in well-trained, thirsty mice (Figure 2Q, left). After these mice were allowed to drink freely until quenched, the same auditory cue no longer elicited anticipatory licking (Figure 2Q, left). We recorded BLA ACh activity in both states and observed a significant reduction in reward CS-evoked ACh activity in quenched mice (Figures 2Q and 2R), consistent with a decrease of salience for the water-predicting cue in quenched mice. Additionally, we observed that BLA ACh responses to shock became significantly larger in the quenched state than in the thirsty state (Figures 2S and 2T), indicating that shock became more salient when homeostatic needs were met.

Taken together, these results demonstrate that BLA ACh activities represent the dynamics of behavioral salience tracked by animals’ motivational engagement across various behavioral contexts or homeostatic states.

### Opposing effects of NAc D1 and D2 MSNs on ACh release in the BLA

We next want to explore how the salience, as represented by BLA ACh activity, is modulated. As previously acknowledged, BLA ACh originates from the basal forebrain, which includes the nucleus basalis of Meynert, diagonal band of Broca, medial septum, and SI ^25,27,28^. It has long been known that basal forebrain cholinergic neurons receive the most prominent inputs from the NAc ^41–44^, but the function of this connection has not yet been explored. We hypothesized that the NAc might regulate BLA ACh levels via its projection onto the basal forebrain cholinergic system. To test this, we injected AAV with Cre-inducible expression of the light-gated cation channel channelrhodopsin-2 (ChR2) into the NAc of *D1-Cre* or *D2-Cre* mice to express ChR2 in two major neuronal populations of the NAc, and injected AAV expressing ACh sensor (ACh3.8m) into the BLA (Figures 3A, 3B, 3H, and 3I). Optical fibers were subsequently implanted in the NAc and BLA for photostimulation and fiber photometry, respectively (Figure S3A). To avoid the laser artifacts to the photometry signal, we interleaved the photometry recording and optogenetic laser stimulation at the frequency of 20 Hz (Figure S3B; see Methods).

**Figure 3.**
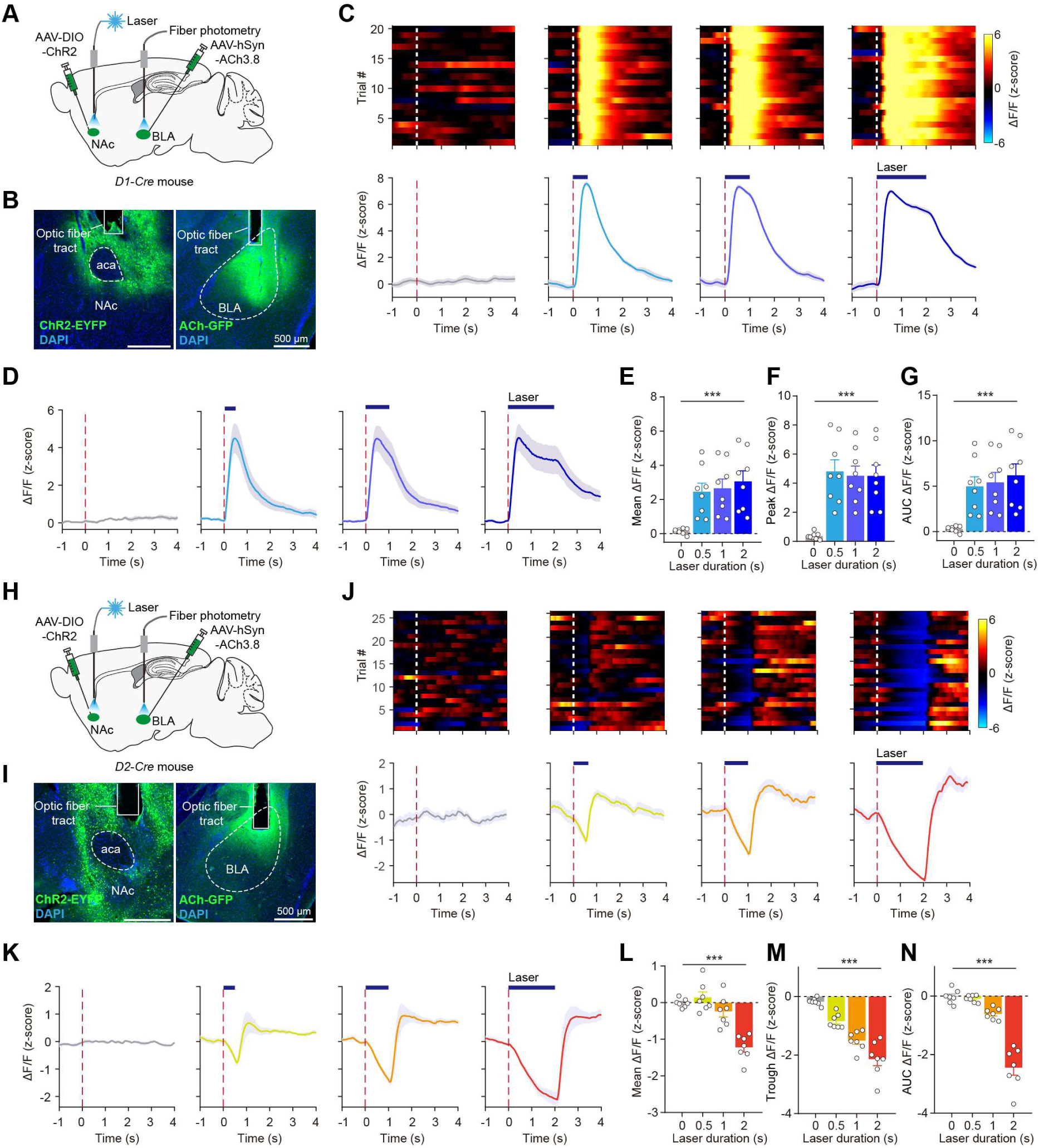
NAc D1 and D2 MSNs opposingly regulate the release of ACh in the BLA. **(A)** Schematic of the strategy to record BLA ACh while activating NAc D1 MSNs. **(B)** Left: representative image of ChR2 expression in NAc D1 MSNs. Right: representative image of ACh3.8 expression in the BLA. **(C)** Trial-by-trial (top) and average (bottom) responses of BLA ACh while activating NAc D1 MSNs with different laser durations in an example session. **(D)** Average BLA ACh responses (n = 8 mice) while activating NAc D1 MSNs with different laser durations. **(E-G)** Quantification of the mean (E) (F_(3, 28)_ = 7.28, p < 0.001), peak (F) (F_(3, 28)_ = 11.02, p < 0.001), and area under curve (AUC) (G) (F_(3, 28)_ = 7.32, p < 0.001) of BLA ACh responses while activating NAc D1 MSNs with different laser durations (n = 8 mice). One-way ANOVA. **(H)** Schematic of the strategy to record BLA ACh while activating NAc D2 MSNs. **(I)** Left: representative image of ChR2 expression in NAc D2 MSNs. Right: representative image of ACh3.8 expression in the BLA. **(J)** Trial-by-trial (top) and average (bottom) responses of BLA ACh while activating NAc D2 MSNs with different laser durations in an example session. **(K)** Average BLA ACh responses (n = 7 mice) while activating NAc D2 MSNs with different laser durations. **(L-N)** Quantification of the mean (L) (F_(3, 24)_ = 22.52, p < 0.001), trough (M) (F_(3, 24)_ = 36.08, p < 0.001), and AUC (N) (F_(3, 24)_ = 22.1, p < 0.001) of BLA ACh responses while activating NAc D2 MSNs with different laser durations (n = 7 mice). One-way ANOVA. ***p < 0.001. Data are presented as mean ± SEM. Shaded areas represent SEM.

To assess whether NAc D1 MSNs regulate BLA ACh, we activated NAc D1 MSNs while simultaneously recording BLA ACh release. We presented the laser stimulation with durations of 0, 0.5, 1.0 or 2.0 seconds interleaved in one session (Figures S3C-S3E). We observed that activation of NAc D1 MSNs induced robust elevation of BLA ACh responses (Figures 3C and S3C). The ACh responses scaled with the duration of laser stimulation, with longer durations resulting in larger area under the curve (AUC) of ACh responses (Figures 3C-3G and S3C). As to the control mice with GFP expression in the NAc, the laser stimulation had no effect on BLA ACh responses (Figures S3E-S3K). These data reveal that activation of NAc D1 MSNs robustly increases BLA ACh levels.

Next, we set out to test whether stimulation of NAc D2 MSNs affects BLA ACh levels. Employing a similar approach to that used with NAc D1 MSNs, we optogenetically activated NAc D2 MSNs and monitored BLA ACh signal with fiber photometry (Figures 3H and 3I). Interestingly, we found that activation of NAc D2 MSNs evoked potent suppression of BLA ACh responses (Figures 3J and S3D). The suppression effect scaled with the duration of laser stimulation, with longer durations inducing a greater suppression effect (Figures 3J-3N). These results show that NAc D2 MSNs potently decreased BLA ACh levels. Together, these data demonstrate that distinct populations of NAc MSNs have opposing effects in modulating ACh release in the BLA.

### NAc MSNs modulate ACh release in the BLA via their projections to the SI

To address what circuit mechanisms account for the accumbal modulation of ACh release, we performed anterograde tracing for NAc D1 and D2 MSNs, and observed dense axonal terminals in the SI (Figures S3L-S3N). Although basal forebrain ChAT neurons project to many target regions, BLA-projecting ChAT neurons are preferentially localized in the SI (Figures S4A-S4C) and the population of BLA-projecting SI ChAT neurons predominantly targets the BLA only (Figures S4D-S4F), suggesting potentially unique anatomical and functional characteristics. As a first step to test whether activation of SI ChAT neurons induces ACh release in the BLA, we expressed ChR2 in SI ChAT neurons and ACh sensor in the BLA (Figures S4G and S4H). We found that brief stimulation of SI ChAT neurons induced robust ACh release in the BLA (Figures S4I-S4K). The enhancement effect scaled with the duration of laser stimulation, with longer durations resulting in a greater enhancement effect (Figures S4K-S4N).

To further examine the causal effect of NAc➔SI pathway on BLA ACh release, we sought to perform optogenetic stimulation of the NAc➔SI pathway while simultaneously recording BLA ACh release. To this end, we infected NAc D1 or D2 MSNs with ChR2, and implanted optical fibers in the SI above the axons originating from the infected neurons, and infected BLA neurons with ACh sensor, and implanted optical fibers in the BLA to detect ACh release (Figures 4A, 4B, 4F, and 4G). Similar to the stimulation of cell bodies of NAc D1 and D2 MSNs, we found robust increase and decrease of BLA ACh responses induced by activation of the NAc^D1^➔SI and NAc^D2^➔SI pathways, respectively (Figures 4C-4E and 4H-4J). These data show that NAc MSNs modulated BLA ACh levels via the projections to the SI.

**Figure 4.**
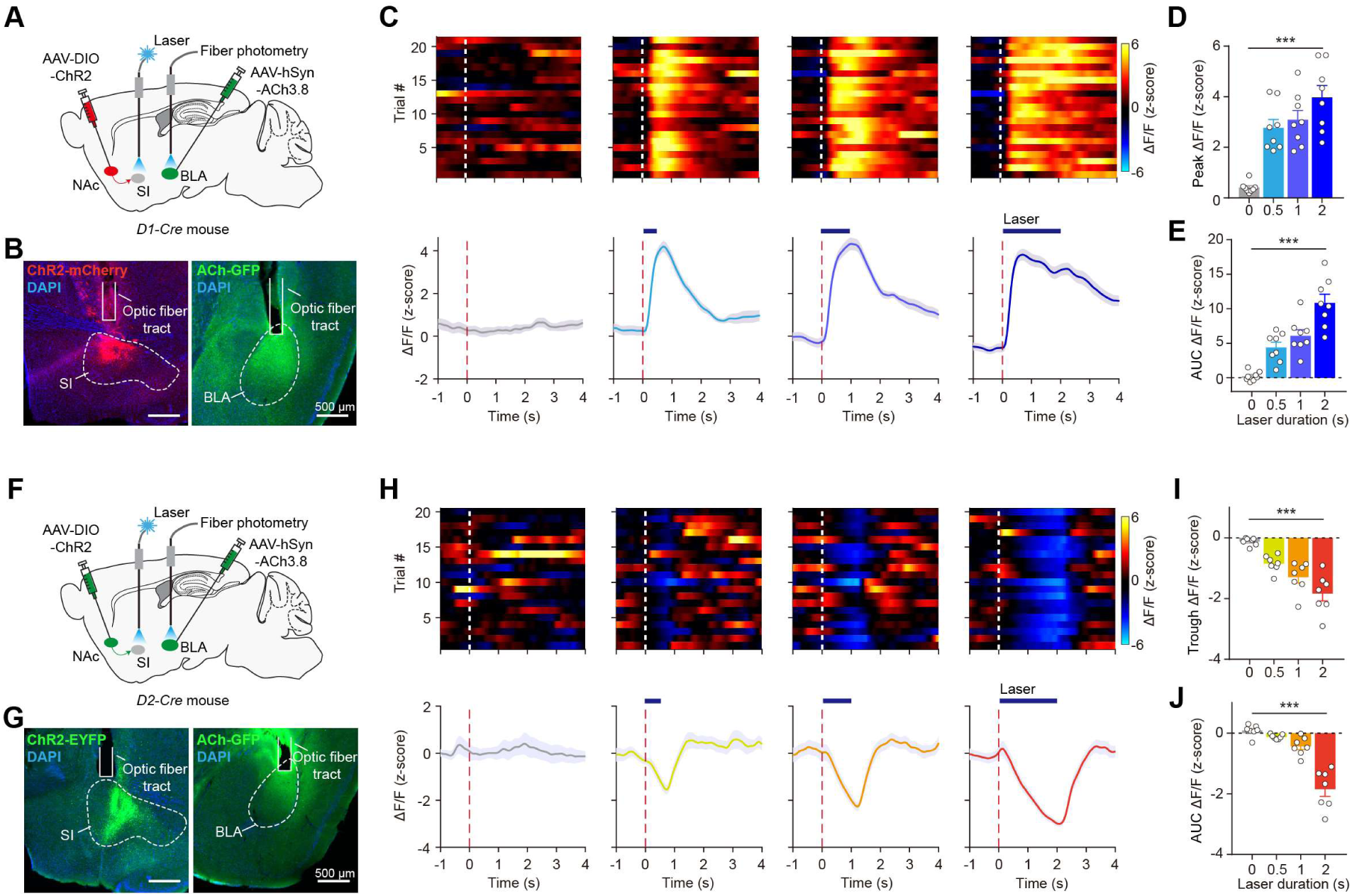
Activation of NAc^D1^➔SI and NAc^D2^➔SI pathways opposingly regulates the release of ACh in the BLA. **(A)** Schematic of the strategy to record BLA ACh while activating NAc^D1^➔SI. **(B)** Left: representative image of ChR2-expressing NAc^D1^ projections to the SI. Right: representative image of ACh3.8 expression in the BLA. **(C)** Trial-by-trial (top) and average (bottom) responses of BLA ACh while activating NAc^D1^➔SI pathway with different laser durations in an example session. **(D, E)** Quantification of the peak (D) (F_(3, 28)_ = 31.54, p < 0.001) and AUC (E) (F_(3, 28)_ = 22.5, p < 0.001) of BLA ACh responses while activating NAc^D1^➔SI pathway with different laser durations (n = 8 mice). One-way ANOVA. **(F)** Schematic of the strategy to record BLA ACh while activating NAc^D2^➔SI. **(G)** Left: representative image of ChR2-expressing NAc^D2^ projections to the SI. Right: representative image of ACh sensor expression in the BLA. **(H)** Trial-by-trial (top) and average (bottom) responses of BLA ACh while activating NAc^D2^➔SI pathway with different laser durations in an example session. **(I, J)** Quantification of the trough (I) (F_(3, 24)_ = 19.05, p < 0.001) and AUC (J) (F_(3, 24)_ = 22.44, p < 0.001) of BLA ACh responses while activating NAc^D2^➔SI pathway with different laser durations (n = 7 mice). One-way ANOVA. ***p < 0.001. Data are presented as mean ± SEM. Shaded areas represent SEM.

### Activation of SI cholinergic neurons and SI^ChAT^➔BLA pathway does not affect valence

Further, we aim to test whether stimulating SI ChAT neurons would drive any form of preference or aversion. To achieve this, we expressed ChR2 selectively into SI ChAT neurons, and positioned the optic fibers above the SI region for laser delivery (Figure S5A). To assess the effects, we subjected these *ChAT-Cre* mice to a real-time place preference test, wherein photo-activation of ChAT neurons was contingent on their entry into a particular side of a chamber (Figure S5B). We found that activation of SI ChAT neurons did not prompt any preference for a specific location (Figure S5C), and it had no impact on the movement speed (Figure S5D) or distance covered by the mice (Figure S5E). Furthermore, we explored whether optogenetic stimulation of the SI^ChAT^➔BLA pathway would induce any observable valence-related behaviors (Figure S5F). Consistent with prior observations ^15,25^, we found that activation of the SI^ChAT^➔BLA pathway did not induce any place preference (Figures S5G and S5H), or affect the movement (Figures S5I and S5J). In addition, activation of this pathway failed to support self-stimulation (Figures S5K-S5N). Taken together, these observations provide evidence that BLA ACh does not actively drive valence-related responses.

### Activation of NAc→SI pathway had no effect on cortical and hippocampal ACh release

To ask whether the stimulation effect of the NAc→SI pathway is selective to ACh release in the BLA, we assessed ACh release in other brain regions originating from the basal forebrain. In addition to the BLA, SI ChAT neurons also project to various other target regions, including the cerebral cortex and hippocampus (Figures 5A-5C). Specifically, we chose to optogenetically manipulate the NAc→SI pathway and monitor the ACh signal in cortical regions, including the medial prefrontal cortex (mPFC) and auditory cortex (AUD), which have been extensively characterized for ACh activity, as well as in the CA1 region of the hippocampus ^27^. We expressed ChR2 in the NAc D1 or D2 MSNs, and implanted optic fibers in the SI for pathway stimulation, and expressed ACh sensor in the mPFC, the AUD, and the CA1 for ACh monitoring (Figures 5D, 5G, and 5J). Both reward and punishment induced a robust increase of ACh release in the mPFC, the AUD, and the CA1 (Figure S6). However, in contrast to the effect observed in the BLA, the NAc^D1^➔SI stimulation did not evoke an immediate and potent increase of ACh signal in the mPFC, the AUD, and the CA1 (Figures 5E, 5H, 5K, and 5M; S7A, S7C, S7E, S7G, S7I, and S7K). The amplitude of ACh responses in the cortex and hippocampus was quite small, compared to that of the BLA (Figure 5M). Although there were some detectable changes in the mPFC, AUD, and the CA1, the peak time lagged the laser stimulation significantly (Figure 5N), suggesting an indirect effect on these areas. Activation of NAc^D2^➔SI pathway did not induce any changes to ACh release in the mPFC, the AUD, and the CA1 (Figures 5F, 5I, 5L, and 5O; S7B, S7D, S7F, S7H, S7J, and S7L). Notably, the suppression effect is only present in the BLA for activating NAc D2 MSNs (Figure 5O), highlighting a unique role of D2 MSNs in modulating BLA ACh levels. These results demonstrate that the modulatory effect of NAc→SI pathway is specific to the BLA.

**Figure 5.**
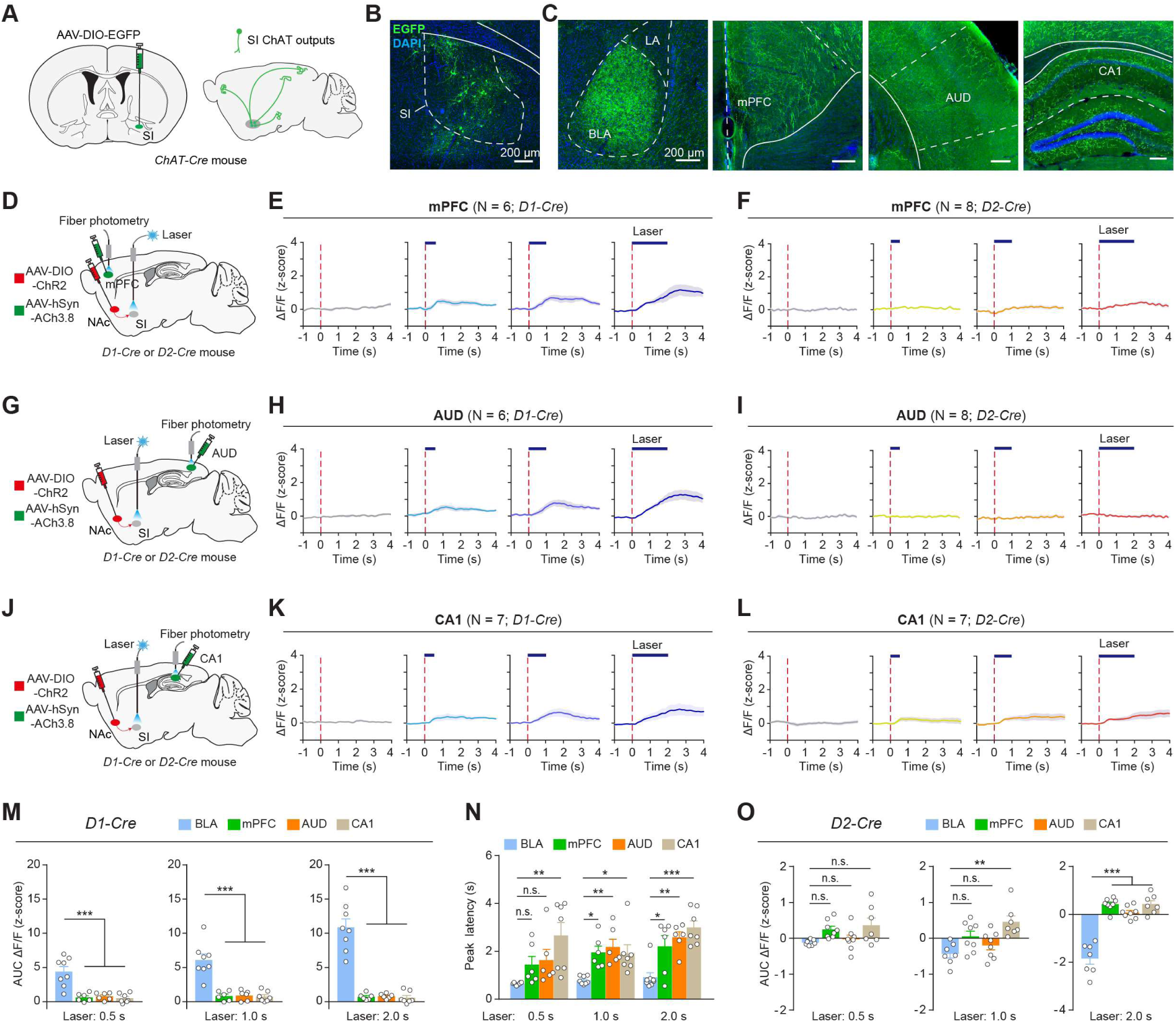
Effects of NAc➔SI activation on the ACh release in the cortex and hippocampus. **(A)** Schematic of anterograde tracing of SI ChAT neurons. **(B, C)** Example coronal sections showing EGFP-expressing cell bodies in the SI (B) and their labeled axonal downstream in the BLA, mPFC, AUD, and CA1 (C). **(D)** Schematic of the strategy to record mPFC ACh while activating NAc^D1^➔SI or NAc^D2^➔SI. **(E, F)** Average mPFC ACh responses across mice while activating NAc^D1^➔SI (E) and NAc^D2^➔SI (F) with different laser durations. **(G)** Schematic of the strategy to record AUD ACh while activating the NAc^D1^➔SI or NAc^D2^➔SI pathway. **(H, I)** Average AUD ACh responses across mice while activating the NAc^D1^➔SI (H) and NAc^D2^➔SI (I) with different laser durations. **(J)** Schematic of the strategy to record CA1 ACh while activating NAc^D1^➔SI or NAc^D2^➔SI. **(K, L)** Average CA1 ACh responses across mice while activating NAc^D1^➔SI (K) and NAc^D2^➔SI (L) with different laser durations. **(M)** Quantification of the ACh release in the mPFC, AUD, and CA1 while activating NAc^D1^➔SI in comparison to ACh release in the BLA (left, F_(3, 23)_ = 16.58, p < 0.001; middle, F_(3, 23)_ = 22.96, p < 0.001; right, F_(3, 23)_ = 47.73, p < 0.001; one-way ANOVA followed by Tukey’s test). **(N)** Quantification of peak latency of ACh signal in the BLA, mPFC, AUD, and CA1 while activating NAc^D1^➔SI (left, F_(3, 23)_ = 5.36, p = 0.006; middle, F_(3, 23)_ = 6.08, p= 0.003; right, F_(3, 23)_ = 10.86, p < 0.001; one-way ANOVA followed by Tukey’s test). **(O)** Quantification of the ACh release in the BLA, mPFC, AUD, and CA1 while activating NAc^D2^➔SI in comparison to ACh release in the BLA (left, F_(3, 26)_ = 3.36, p = 0.034; middle, F_(3, 26)_ = 6.89, p = 0.0014; right, F_(3, 26)_ = 50.67, p < 0.001; one-way ANOVA followed by Tukey’s test). *p < 0.05, **p < 0.01, ***p < 0.001; n.s., non-significant (p > 0.05). Data are presented as mean ± SEM. Shaded areas represent SEM.

### SI ChAT neurons form direct connection preferentially with NAc D2, but not D1 MSNs

As distinct subpopulations of NAc MSNs have opposing modulatory effects on the BLA ACh levels, we next sought to understand whether and how NAc D1 and D2 MSNs are connected differentially to SI ChAT neurons. Given that both NAc D1 and D2 MSNs are GABAergic and release inhibitory neurotransmitters, it is unlikely for the majority of D1 MSNs to form direct monosynaptic connections with SI ChAT neurons. Conversely, D2 MSNs are more likely to have direct monosynaptic connections with these neurons. We proposed that NAc D1 MSNs could activate SI ChAT neurons by inhibiting intermediate, tonically active GABAergic afferents to ChAT neurons, while D2 MSNs directly inhibit ChAT neurons.

To test this prediction, we performed high-resolution volumetric imaging of NAc axon terminals and ChAT cell bodies to directly visualize the connections between NAc MSNs and SI ChAT neurons. By injecting an AAV expressing a membrane-targeted green fluorescent protein (mGFP) and synaptophysin, which is found on nearly all presynaptic vesicles, fused with a red fluorescent protein, mRuby, in a Cre-dependent manner into the NAc of *D1-Cre* or *D2-Cre* mice (Figures 6A and 6D), we could identify the synaptic targets of NAc MSNs. This method confines mRuby expression to pre-synaptic terminals, enabling us to differentiate between potential synaptic targets and passing axons. We observed robust labeling of synaptic terminals originating from NAc MSNs in the SI (Figures 6B, 6C, 6E, and 6F). To quantify the number of synaptic connections, we counted the red spots present within 1.0 μm distance from ChAT neurons, which were immunostained with green fluorescence (Figure 6G). We found that the number of synaptophysin-mRuby spots around ChAT neurons for D2 MSNs was significantly higher than that for D1 MSNs (Figure 6H), indicating that D2 MSNs predominantly target SI ChAT neurons, thereby inhibiting their Ach release.

**Figure 6.**
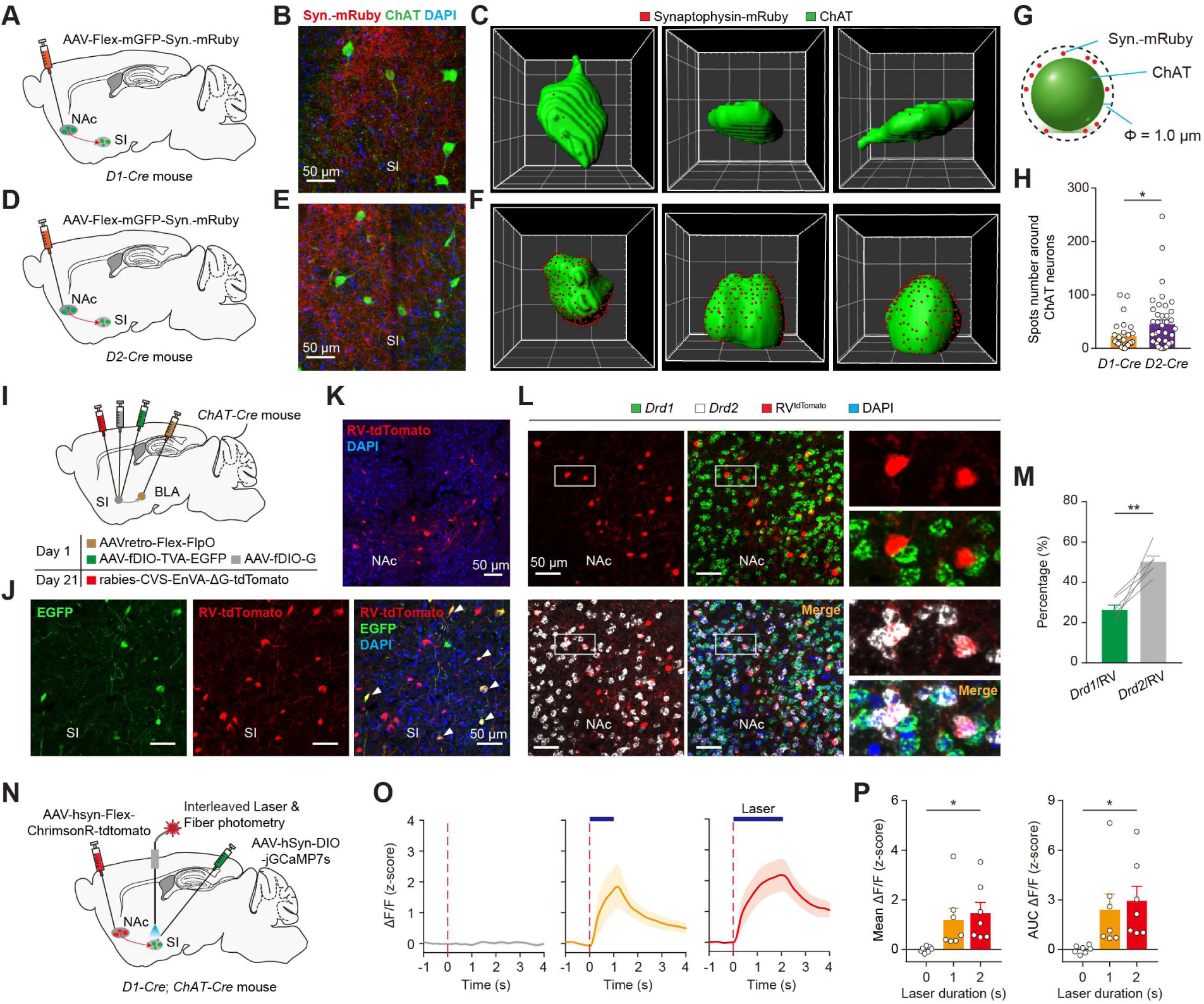
Synaptic and functional connections between NAc MSNs and SI ChAT neurons. **(A, D)** Schematic of the approach. **(B, E)** Confocal images showing synaptic puncta (red) of NAc D1 (B) and D2 (E) MSNs and cell body of ChAT neurons (green). **(C, F)** High-magnification images showing 3D reconstructed ChAT neurons and surrounding synaptophysin spots from a representative *D1-Cre* (C) and *D2-Cre* (F) mouse. **(G)** Schematic of synaptophysin spot quantification around ChAT neurons. **(H)** Quantification of synaptophysin spots surrounding ChAT neurons for D1 and D2 MSNs (p = 0.032, t-test). **(I)** Schematic of the approach. **(J)** Confocal images from a mouse prepared as in (I), showing the BLA-projecting ChAT neurons in the SI infected by the helper viruses (green) and the cells infected by the rabies virus (RV) (red). The starter cells are yellow, as indicated by the arrowhead. **(K)** Representative image of the NAc, showing input neurons labeled by the RV. **(L)** Confocal images of in situ hybridization for RV-tdTomato, Drd1, and Drd2 in the NAc. Right, high-magnification views of the boxed area on the left. **(M)** Quantification of the percentage of D1 or D2 NAc MSNs among RV-tdTomato cells (n = 6; p = 0.0029, paired t-test). **(N)** Schematic of the approach. **(O)** Average SI ChAT neural activities while activating the NAc^D1^➔SI pathway with different laser durations (n = 7 mice). **(P)** Quantification of the mean (left; F_(2, 18)_ = 4.35, p = 0.03) and AUC (right; F_(2, 18)_ = 4.29, p = 0.03) of SI ChAT neural activities (n = 7 mice). One-way ANOVA. *p < 0.05, **p < 0.01. Data are presented as mean ± SEM.

To confirm that BLA-projecting SI ChAT neurons indeed receive inputs from the NAc, we utilized an intersectional monosynaptic retrograde rabies system, which was injected into the SI of ChAT-Cre mice in a pathway- and cell-type-specific manner (Figures 6I and 6J). We found that BLA-projecting SI ChAT neurons received substantial inputs from the NAc (Figure 6K). To further characterize the specific cell types of NAc MSNs with monosynaptic inputs onto SI ChAT neurons, we performed single-molecule *in situ* hybridization on retrogradely labeled NAc MSNs infected by rabies virus (Figure 6L). We found that the vast majority of rabies virus-labeled cells expressed the dopamine receptor gene *Drd2*, but not *Drd1* (Figure 6M). This data shows that NAc D2 MSNs directly targeted SI ChAT neurons, which could inhibit ACh release in the BLA. In the basal forebrain, the vast majority of neurons are cholinergic and GABAergic ^45^. ChAT neurons receive inputs from local GABAergic neurons ^43^ and can be inhibited by local GABAergic neurons ^17^. To further test whether NAc D1 MSNs could disinhibit SI ChAT neurons, we recorded SI ChAT neural activities while activating the NAc^D1^➔SI pathway (Figure 6N). We observed that activation of the NAc^D1^➔SI pathway resulted in a potent increase in SI ChAT neural activities (Figures 6O and 6P). Our results thus support the notion that NAc D1 MSNs inhibit local SI GABAergic neurons, resulting in disinhibition of ChAT neurons, hence increased ACh release in the BLA.

### Activation of NAc^D1^➔SI facilitates and NAc^D2^➔SI delays associative learning

Learning how environmental stimuli predict the availability of food and other natural rewards is critical for survival. Salient stimuli capture attention, facilitating associative learning. If the NAc modulates the salience representation of BLA ACh, then activation of the NAc➔SI pathway, which either enhances or suppresses BLA ACh levels, should affect the efficiency of cue-reward learning. To this end, we trained mice to perform a cue-reward learning task as described before ^25^. In this task, water-restricted mice were trained to perform a correct nose poke when signaled by an auditory cue to receive a water reward on a variable intertrial interval (Figures 7A and 7B). We first characterized the *in vivo* activity of BLA ACh in this task (Figure S8). We observed that BLA ACh release occurred in response to various events across the learning of the task (Figures S8A and S8B). In rewarded trials of the pre-training stage, the highest levels of ACh release occurred immediately after the reward retrieval (entry into the reward port) (Figures S8A and S8B). As the training began, ACh release during reward trials showed a marked shift toward the time before nose poking reward port (Figures S8A and S8B). As animals became proficient in the task, there was a significant increase in the magnitude and a significant decrease in the peak latency of ACh release upon cue presentation (Figures S8C-S8F). This indicates that as animals learned the cue-reward contingency, the cue became a more salient event. At this time point, there was still a peak of ACh release at the time of reward, but its magnitude was diminished, and the peak occurred earlier (Figure S8G). Consistent with previous findings ^25^, the ACh responses shifted as mice gradually learned the task. This shows that BLA ACh is actively involved in the reward learning association.

**Figure 7.**
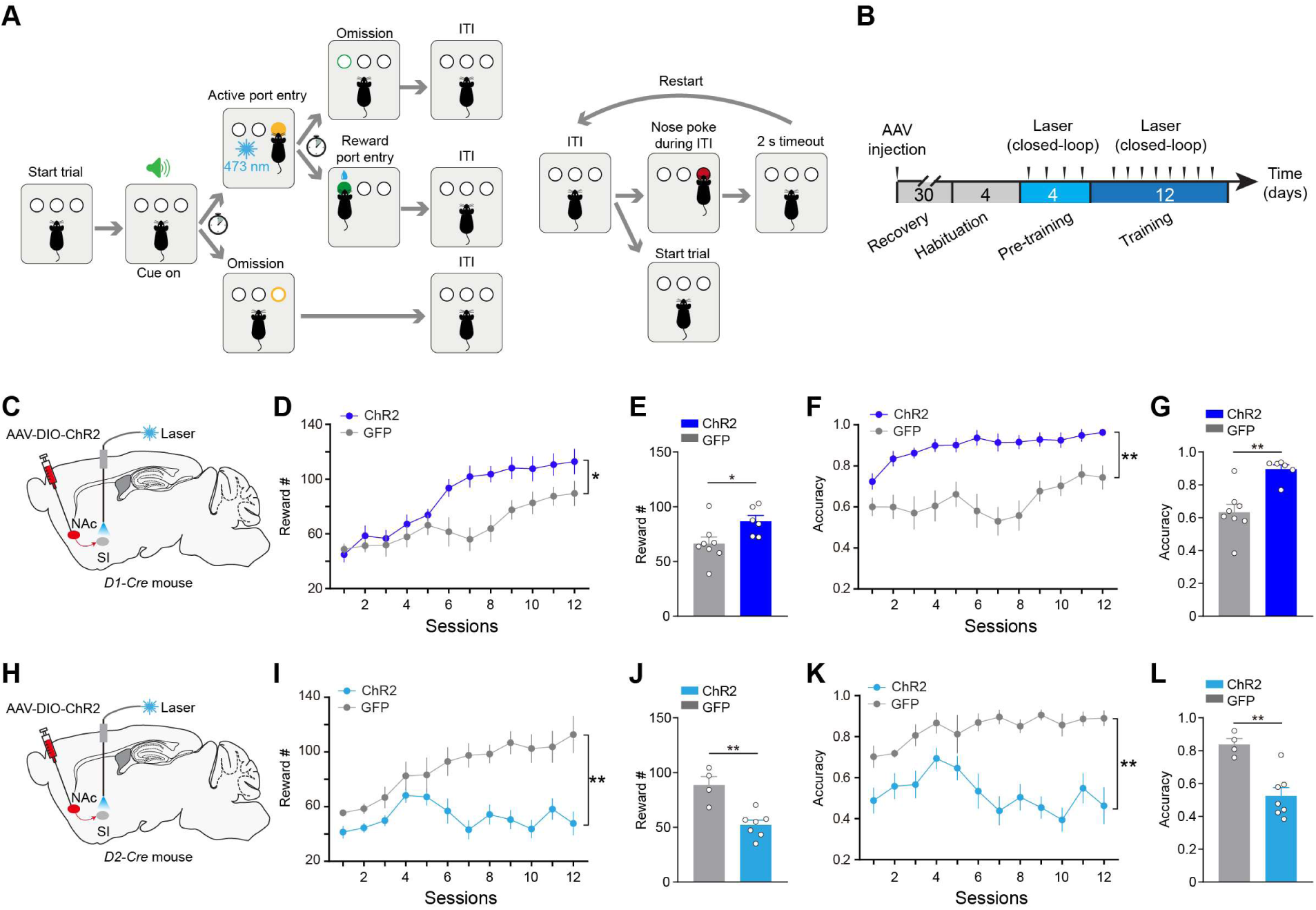
Activation of NAc^D1^➔SI and NAc^D2^➔SI facilitates and impairs reward associative learning, respectively. **(A)** Schematics of the cue-reward learning task. In the pre-training phase, there was no consequence for incorrect nose pokes (left). In training, timeout punishment was triggered by incorrect nose pokes (right). **(B)** Schematic of the experimental procedure. **(C)** Schematic of the strategy to activate NAc^D1^➔SI pathway. **(D, E)** Rewards earned by the ChR2 (n = 6) and GFP (n = 8) mice in each session (D) (interaction, F_(11, 132)_ = 4.18, p < 0.0001; group, F_(1, 12)_ = 5.75, p = 0.034; two-way ANOVA), and average across sessions (E) (p = 0.034, t-test) with closed-loop stimulation of NAc^D1^➔SI in the cue-reward learning task. **(F, G)** Behavioral accuracy of the ChR2 (n = 6) and GFP (n = 8) mice in each session (F) (interaction, F_(11, 132)_ = 2.01, p = 0.032; group, F_(1, 12)_ = 18.32, p = 0.0011; two-way ANOVA), and average across sessions (G) (p = 0. 0011, t-test) with closed-loop stimulation of NAc^D1^➔SI in the cue-reward learning task. **(H)** Schematics of the strategy to activate NAc^D2^➔SI pathway. **(I, J)** Rewards earned by the ChR2 (n = 7) and GFP (n = 4) mice in each session (I) (interaction, F_(11, 99)_ = 6.25, P < 0.001; group, F_(1, 9)_ = 19.52, p = 0.0017; two-way ANOVA), and average across sessions (J) (p = 0.0017, t-test) with closed-loop stimulation of NAc^D2^➔SI in the cue-reward learning task. **(K, L)** Behavioral accuracy of the ChR2 (n = 7) and GFP (n = 4) mice in each session (K) (interaction, F_(11, 99)_ = 3.16, p = 0.0011; group, F_(1, 9)_ = 18.21, p = 0.0021; two-way ANOVA), and average across sessions (L) (p = 0.0021, t-test) with closed-loop stimulation of NAc^D2^➔SI in the cue-reward learning task. *p < 0.05, **p < 0.01. Data are presented as mean ± SEM.

Given the observed shift of BLA ACh towards tone onset during acquisition of cue-reward learning, we hypothesized that BLA ACh may potentiate learning the cue-reward contingency. To test this, we optogenetically activated the NAc^D1^➔SI pathway in a closed-loop manner, contingent on the correct nose poke and reward retrieval, thereby boosting BLA ACh release during behavior. Activation of NAc^D1^➔SI pathway significantly facilitated animals’ learning rate (Figures 7C-7G). Conversely, activation of NAc^D2^➔SI pathway significantly delayed mice’s reward learning (Figures 7H-7L). Since reaction time tends to be shorter when stimuli are more salient ^46,47^, we used reaction time (from cue onset to port entry) as an indicator of stimulus salience, measured during periods without laser stimulation. We found that reaction time was significantly shorter in *D1-Cre* mice compared to controls (Figures S9A-S9C), while it was notably prolonged in *D2-Cre* mice (Figures S9F-S9H). This data suggests that activation of NAc^D1^➔SI pathway promotes salience, leading to quicker responses in mice, whereas NAc^D2^➔SI pathway suppresses salience, resulting in slower responses. Considering the inhibitory role of D2 MSNs in the dorsal striatum on movement, we speculated whether impaired reward learning could be attributed to movement deficits. However, our results indicated that activation of NAc D2 MSNs did not significantly affect general poking behavior (Figures S9I and S9J). Together, our results reveal that distinct NAc neural populations opposingly modulate the BLA ACh levels, leading to opposite changes in the learning rate in a cue-outcome contingency setting.

## DISCUSSION

Previous studies of selective attention and the salience of behaviorally relevant stimuli have primarily focused on the frontoparietal cortex and the thalamus ^1,38,48,49^. Here we have identified the NAc, located within the ventral striatum of the basal ganglia, as a key regulator of salience assignment. Specifically, we demonstrate that two distinct subpopulations of striatal MSNs exert opposing control over salience signals via modulating SI ChAT neurons (Figures 3 and 4). The NAc is uniquely positioned at the crossroads of emotional, cognitive, and motor circuits, enabling the integration of diverse information ^50,51^. While D1 and D2 MSNs in the dorsal striatum are segregated based on their respective projection targets, NAc D1 and D2 MSNs project to several overlapping downstream targets, including the SI (Figures 4B, 4G, and S3L-S3N). Then how do NAc D1 and D2 MSNs exhibit opposing roles in salience assignment? Given the diverse cortical and limbic inputs received by the NAc ^52–54^, future investigations are warranted to delineate the contributions of individual inputs while examining salience control by NAc MSNs.

Salience, defined as the absolute (i.e., unsigned) value of a stimulus, is a concept often intertwined with valence, making their neural separation challenging. Activation of NAc D1 or D2 MSNs can induce potent valence-related responses, either rewarding or aversive ^54–57^. In contrast, stimulation of SI ChAT neurons or the SI➔BLA pathway failed to evoke any valence-specific responses (Figure S5), consistent with ACh’s representation of salience rather than valence. This suggests that the rewarding and aversive responses may be mediated by other downstream targets of the NAc or the SI, such as the lateral hypothalamus and ventral tegmental area ^58–61^. Our results highlight the ability to disentangle salience from valence at the neural circuit level.

Associative learning, a fundamental cognitive process that allows organisms to extract relationships among environmental cues, actions, and their outcomes, underlies the formation of memories and prediction of future events ^49,62^. To achieve this, our brains need to not only form and store these associations but also assign importance or “salience” to relevant cues in the environment, thus facilitating learning and memory processes. Activation of the cholinergic system is capable of modulating synaptic plasticity by altering synaptic strength and regulating secretion of neurotrophic factors ^14,63–66^. Through these mechanisms, BLA-projecting cholinergic neurons may boost neural plasticity and thereby contribute to associative learning. In line with this notion, we found that enhancing and suppressing ACh release in the BLA via manipulation of NAc D1 and D2 MSNs facilitated and impaired reward associative learning, respectively (Figure 7). It is widely acknowledged that the NAc is involved in various forms of learning and motivational control. In particular, our results reveal the striatal involvement, mediated by NAc➔SI pathway, in the salience assignment as an important contributing aspect of learning and motivational control.

In the striatum, the interaction among local ChAT interneurons, dopaminergic inputs, and MSNs has been extensively explored ^67–69^. In the striatum, axons of midbrain dopamine neurons express high levels of nicotinic ACh receptors ^67^. Activation of these ACh receptors can directly trigger action potentials in striatal dopaminergic axons, resulting in dopamine release ^67,70,71^. Here we showed that the striatal neurons interact with the extra-striatum cholinergic system, the basal forebrain cholinergic system. The NAc MSNs modulated the SI cholinergic system potently, with NAc D1 and D2 MSNs facilitating and suppressing ACh release in the BLA, respectively (Figures 3 and 4). Interestingly, dopamine in the BLA has recently been demonstrated to encode salience ^72^. Future studies will need to determine whether dopamine signals in the BLA are triggered by the cholinergic inputs from the basal forebrain, and whether dopamine and ACh signals both participate in salience assignment in a coordinated manner.

## ACKNOWLEDGEMENTS

We thank M.M. Poo, X. Jin, B. Li, B. Hangya, A. Laszlo, N.L. Xu, and Y.G. Sun for comments on earlier versions of the manuscript, and members of the Xiao laboratory for helpful discussions. We thank Yulong Li for providing us with the ACh3.8 sensor. We thank Qian Hu (the Optical Imaging Core at the CEBSIT) for providing assistance with the analysis of volumetric imaging data. This work was supported by grants from the National Natural Science Foundation of China (32371060 to X.X., 32271065 to H.D.), the Lingang Laboratory (LG-QS-202203-06 to X.X., LG-QS-202203-02 to H.D.), the Chinese Academy of Sciences (to X.X.), and the Benyuan Charity Foundation (to H.D.).

## AUTHOR CONTRIBUTIONS

A.C., H.D., and X.X. designed the study. A.C. conducted most of the experiments with help from Y.L. and H.Z. A.C. analyzed data and prepared figures. H.G. helped with fiber photometry and histology. Y.W. helped with the behavioral training. W.G., Q.S., B.L., Q.Y., and F.X. provided critical conceptual consultation and experimental suggestions. X.X., A.C., and H.D. wrote the paper with inputs from all authors. X.X. supervised all aspects of the work.

## RESOURCE AVAILABILITY

### Lead Contact

Further information and requests for resources and reagents should be directed to and will be fulfilled by the Lead Contact, Xiong Xiao.

### Data and code availability

The authors declare that the data supporting the findings of this study are available within the paper and its supplementary information files. The custom codes that support the findings from this study will be made available online.

## EXPERIMENTAL MODEL AND SUBJECT DETAILS

All experiments were approved by the Animal Care and Use Committee of the Institute of Neuroscience, Chinese Academy of Sciences (Center for Excellence in Brain Science and Intelligence Technology, Chinese Academy of Sciences). We obtained *ChAT-Cre* (Stock No: 006410) mice from Jackson Laboratory. The *D1-Cre* (MMRRC_030989-UCD) and *D2-Cre* (MMRRC_032108-UCD) mice were provided by Dr. Ninglong Xu at the CEBSIT. Male and female mice (2-4 months old) were used for all the experiments. Mice were housed under a 12-h light/dark cycle (7 a.m. to 7 p.m. light) in groups of 2-6 animals, with food and water available *ad libitum* before being used for experiments in the animal facility at the Institute of Neuroscience. Littermates were randomly assigned to different groups before experiments.

## METHOD DETAILS

### Immunohistochemistry

Immunohistochemistry experiments were conducted following standard procedures ^32,73,74^. Briefly, mice were anesthetized with isoflurane and perfused with PBS, followed by 4% paraformaldehyde (PFA). Brains were extracted and post-fixed overnight in 4% PFA at 4 °C. Coronal sections (50- μm) were cut using a vibratome (VT1200S, Leica). Sections were first washed in PBS (5 min), then sections were blocked in 5% normal goat serum or BSA in PBST (0.1% Triton X-100 in PBS) for 1 h at room temperature (RT) and then incubated with primary antibodies overnight at 4 °C. Sections were washed with PBS (3 ×15 min) and incubated with fluorescent secondary antibodies at RT for 2 h. Next, sections were washed twice in PBS, and then incubated with DAPI (Sigma, Cat. No. d9542; dilution, 1:5000) for 10 min. After that, sections were mounted onto slides with Fluoromount-G (Southern Biotech). Images were captured by a 10× objective fluorescent microscope (Olympus, VS120).

The primary antibodies used were: chicken anti-GFP (Abcam, Cat. No. ab13970; dilution, 1:1000), and goat anti-ChAT (Millipore, Cat. No. AB144P; dilution, 1:500). Appropriate fluorophore-conjugated secondary antibodies (Jackson ImmunoResearch Laboratories) were used depending on the desired fluorescence colors.

### Stereotaxic surgery

All surgery was performed under aseptic conditions. Standard surgical procedures were used for stereotaxic injection and implantation, as previously described ^32,73,74^. Briefly, mice were anesthetized with isoflurane (1-2% in a mixture with oxygen), and head-fixed in a stereotaxic injection apparatus (Rotational Digital Stereotaxic Frame, RWD Life Science, Shenzhen, China). For virus injection, the following stereotaxic coordinates were used for the NAc: 1.40 mm anterior from bregma, 1.50 mm lateral from the midline, 4.60 mm vertical from skull surface; for the SI, 0.00 mm from bregma, 1.65 mm lateral from the midline, 5.10 mm vertical from skull surface; for the BLA, 1.50 mm posterior from bregma, 3.40 mm lateral from the midline, 4.80 mm vertical from skull surface; for the mPFC, 1.80 mm anterior from bregma, 0.50 mm lateral from the midline, 2.10 mm from skull surface; for the AUD, 2.10 mm posterior from bregma, 3.80 mm from the midline, 2.80 mm from skull surface; for the CA1, 2.00 mm posterior from bregma, 1.40 mm from the midline, 1.50 mm from skull surface.

For fiber photometry recordings, optic fibers (core diameter, 200 mm; N.A., 0.5; Inper, Hangzhou, China) were implanted 50 μm above the coordinates for viral injection. For the optogenetic experiments, optic fibers (core diameter, 200 mm; N.A., 0.37; Inper, Hangzhou, China) were implanted 200 μm above the coordinates for viral injection. Then a metal head-bar (for head-restraint) was subsequently mounted onto the skull with dental cement. We waited for a minimum of 3 weeks before starting the experiments with these mice.

### Viral vectors

The following adeno-associated viruses (AAVs) were obtained from Shanghai Taitool Bioscience (Shanghai, China): AAV2/8-hSyn-EGFP-WPRE-pA, AAV2/8-hEF1α-DIO-hChR2-EYFP-WPRE-pA, AAV2/8-hEF1α-DIO-hChR2-mCherry-WPRE-pA, AAV2/2Retro-Plus-hEF1a-DIO- EGFP-WPRE-pA, AAV2/2Retro-CAG-FLEX-FlpO-WPRE-pA. The AAV2/8-EF1α-DIO-ChR2-mCherry, AAV2/9-EF1α-DIO-EGFP, AAV-nEF1α-fDIO-EGFP, AAV2/9-hsyn-Flex-mGFP-T2A-Synaptophysin-mRuby and AAV2/8-hSyn-ACh3.8 were produced by BrainCase (Shenzhen, China). The components of the rabies viral tracing system were also produced by BrainCase: rAAV-nEF1α-fDIO-EGFP-T2A-TVA, rAAV-nEF1α-fDIO-oRVG, AAV2/9-EF1α-DIO-EGFP-T2A-TVA, AAV2/9-EF1α-DIO-N2c-G, RV-CVS-EnVA-ΔG-tdTomato (the modified rabies virus). All viral vectors were aliquoted and stored at −80°C until use.

### Mapping monosynaptic inputs with pseudotyped rabies virus

To map monosynaptic inputs onto ChAT^SI➔BLA^ neurons, we implemented a pathway-specific and cell-specific tracing strategy utilizing a refined rabies virus system, as previously detailed ^32,74^. We injected the BLA of *ChAT-Cre* mice with the AAVretro-Flex-FlpO, which allows selective expression of FlpO within ChAT^SI➔BLA^ neurons. Subsequently, in the same surgery, we injected the SI of these mice with a mixture of AAV-fDIO-TVA-EGFP and AAV-fDIO-oG (1:2 in volume, for a total of 0.1∼0.2 µl) that express the following components in a Flp-dependent manner: a fluorescent reporter EGFP, TVA (which is a receptor for the avian virus envelope protein EnvA), and the rabies envelope glycoprotein (oG). Three weeks after the first injection, mice were injected in the SI with RV-CVS-EnVA-ΔG-tdTomato (0.1 µl), a rabies virus that is pseudotyped with EnvA, lacks the envelope glycoprotein, and expresses tdTomato. This variant of the rabies virus has been demonstrated to exhibit enhanced retrograde trans-synaptic transfer and reduced neurotoxicity. Histological examination of brain tissue was conducted one week post-rabies virus administration.

### Fluorescent *in situ* hybridization

Single molecule fluorescent *in situ* hybridization was employed to detect the expression of *Drd1* and *Drd2* mRNAs in the nucleus accumbens (NAc) of adult *ChAT-Cre* mice following rabies virus injection, with tdTomato expression in NAc neurons projecting to the substantia innominata (SI). For tissue preparation, mice were anesthetized with isoflurane and perfused with DEPC-PBS and 4% PFA. Brains were taken out and fixed overnight at 4 °C and dehydrated by 30% sucrose in DPEC-PBS. Brains were sectioned at 20 μm and sections containing the NAc were collected and stored at −80°C until processing. Hybridization was carried out using the RNAscope kit (ACDBio).

On the day of the experiment, sections were washed once in RNA-free H2O (1 min), and dehydrated using 100% ethanol (twice, 3 min each). Sections were then dried at RT and incubated with Hydrogen Peroxide for 20 min. Retrieval reagents were then applied to the sections for 5-7 min, followed by a wash with 100% ethanol (twice, 3 min each) and incubation with Protease Ⅲ for 30 min at RT. Sections were washed in PBS for 5 min at RT before hybridization. Probes against *Drd1* (Cat. No. #406491, dilution 1:50) and *Drd2* (Cat. No. #406501, dilution 1:50) were applied to NAc sections, followed by hybridization at 40°C for 2 hours. After hybridization, sections were washed twice in wash buffer (2 min each) at RT, then incubated with three consecutive rounds of amplification reagents (30 min, 30 min and 15 min at 40°C). Finally, fluorescence detection was carried out by HRP-TSA system. To visualize tdTomato-expressing (rabies virus-infected) cells, we performed immuno-staining using primary antibody rabbit anti-dsRed (Takara Bio, Cat. No. 632496, 1:300) and secondary antibody goat anti-rabbit, Cy3 (Jackson ImmunoResearch, Cat. No. 111-655-003, 1:2,000) after *in situ* hybridization. Finally, sections were incubated with DAPI for 2 min, and then mounted with coverslip using mounting medium. Images were acquired using a 40× objective confocal microscope (Nikon, C2) and visualized and processed using ImageJ.

### Measurement of synaptophysin spots

Synaptic terminals in the SI were labeled using synaptophysin-mRuby, a presynaptic marker. Volumetric images were captured by 40× objective confocal microscopy (FV3000, Olympus). Subsequently, we utilized Imaris 8.1 software (Bitplane) for neuronal cell body and synaptophysin spot identification. During the analysis, we meticulously adjusted the diameter parameter to ensure the accurate detection of ChAT neurons, filtering out those close proximity to the image boundary. To achieve 3D segmentation of ChAT neurons, we utilized the Imaris surface and mask selection feature. Following this step, we employed ImageJ 3D Fast Filters to expand the cellular boundary by 1.0 μm. Synaptophysin spots were detected based on their distance from the surface of ChAT neuron cell body. Only spots located within a maximum distance of 1.0 μm from the surface of ChAT neurons were considered positive and included for further analysis. This procedure was conducted in three-dimensional space to better represent the complex architecture of the synaptic connectivity. Finally, ImageJ 3D Manager was employed to quantify the number of synapse termini spots surrounding individual ChAT neurons.

### Markerless pose tracking

We used DeepLabCut software (v.2.2.0.5) for tracking the eyes of mice during behavior. The training data set included 120 frames from 6 videos across 6 mice. These frames were meticulously selected to encompass a diverse range of eye sizes and positions, ensuring robust model training and generalization. For eye tracking, we strategically placed eight tracking points evenly distributed along the periphery of the eye. This configuration enabled comprehensive tracking of eye movements and provided sufficient spatial resolution for capturing subtle variations in eye size and position. The training of the model was conducted over 1.0 million iterations with default parameters (trained on 95% of labeled frames, initialized with ResNet-50, batch size of 8).

### Real-time place preference or aversion test

Freely moving mice were initially habituated to a two-sided chamber (20 × 42 × 30 cm) for 10 minutes to assess baseline preferences. In the subsequent test sessions, lasting 10 minutes each, one side of the chamber was designated as the photo-stimulation side, while the other served as the non-stimulation side. The order of stimulation sides was counterbalanced across mice. Upon entry into the stimulation side, photo-stimulation was initiated using a 473-nm laser (Shanghai Laser & Optics Century, Shanghai, China), delivering 15-ms pulses at 20 Hz and 10 mW intensity (measured at the tip of optic fibers). Laser stimulation ceased as the mouse exited the stimulation side. The behavior of the mice was videotaped with a CCD camera interfaced with Bonsai software (Bonsai), which was used to control the laser and extract real-time behavioral parameters, including position, time, distance, and velocity.

### Self-stimulation test

Freely moving mice were individually placed in a custom-designed chamber equipped with two distinct ports. One port, termed the active port, was designated to trigger photo-stimulation (15-ms pulses, 20 Hz, 10 mW; λ = 473 nm) for 2 s upon poking. In contrast, poking into the other port, termed the inactive port, did not elicit any photo-stimulation. Mice were allowed to freely explore and poke both ports during two separate 1-hour testing sessions, conducted on consecutive days. The assignment of the active port in each session was systematically counterbalanced across experimental subjects.

### Pavlovian conditioning task

Three weeks after surgery, water-deprived mice were trained on an auditory Pavlovian conditioning task, during which the mice were head restrained using custom-made clamps and head-bars mounted on the skull. Prior to training, each mouse underwent a one-day habituation period to acclimate to the head-restraint setup. During training, each trial began with a conditioned stimulus (CS), which was a 0.5-s sound (12 kHz or 3 kHz), followed by a 1-s delay and then an unconditioned stimulus (US; the outcome). The US was either a water reward (5 µl) or an electric shock (0.02 mA, 100 ms). The electric shock was administered towards the tail of the animals.

Once mice reached a performance level of at least 80% successful trial, mice were subjected to the probabilistic conditioning task. The trials were categorized into three types: predicted US (70%), US omission (15%), and unpredicted US (15%). In predicted US trials, a 0.5-s sound was followed by a predicted reward (water) or punishment (shock). In US omission trials, a 0.5-s sound was presented but without subsequent reward or punishment. In unpredicted US trials, a reward or punishment was delivered without preceding CS introduction. These contingencies were carefully calibrated to keep mice motivated for the task. Different trial types were presented in a pseudorandomized sequence.

Water rewards were dispensed via a metal spout positioned in front of the mice’s mouths, which also functioned as part of a custom “lickometer” circuit. This circuit registered a lick event each time a mouse completed the circuit by licking the spout. Experimental control, including the delivery of CSs and USs, as well as the recording of licking events, was managed by a custom software written in MATLAB (MathWorks, Massachusetts) interfaced with a Bpod State Machine (Sanworks, NY).

### Go/no-go task

Water-deprived mice were trained in an auditory go/no-go task under head restraint. The training started with habituation, during which mice received water rewards by licking the water spout (2 µl for each lick), with no auditory stimuli presented. Once mice reliably licked the spout (2-3 days), they were subjected to the go/no-go training, consisting of both “go” and “no-go” trials. In go trials, a 12-kHz tone (termed the “go cue”, 0.5 s in duration; CSG) was delivered, followed by a 1-s delay (the “response window”). Licking during this window resulted in a water reward (5 μl). Conversely, in no-go trials, a different 3-kHz tone (termed the “no-go cue”, 0.5 s in duration; CSN) was delivered, followed by a response window (1 s). Liking during this period was punished by an air puff blowing to the face. The go and no-go trials were randomly interleaved. For analysis, trials were sorted into go trials and no-go trials. A correct response during a go trial (“hit”) occurred when the mouse successfully licked the spout during the response window and subsequently received the water reward. A correct response during a no-go trial (“correct rejection”) occurred when the mouse successfully refrained from licking during the response window and thus avoided the air puff.

### Active avoidance task

The task was designed to train mice to actively avoid punishment. In avoid trials, a 3-kHz tone (termed the “avoid cue”, 0.5 s in duration; CSA, the same tone as CSN) was presented, followed by a 1-s decision window. If mice lick the spout during the decision window, they would avoid an unpleasant air puff (200 ms) blowing to the face. Otherwise, mice would receive the air puff immediately after the decision window. Animals were trained one session per day, with each session consisting of 100 trials.

### Cue-reward learning task

Freely moving water-deprived mice underwent a cue-reward learning task within a chamber equipped with three ports: left (reward port), middle (inactive port), and right (active port). Nose pokes were detected via infrared beam breaks. In the habituation phase of the task, mice could receive a reward with a nose poke in either of the ports. During the Pre-training phase, the trial starts with a 6-s auditory cue (the “response window”), mice need to poke the active port during the 6-second window. Upon successful poking, a 4-s reward collection window was followed, indicated by an illuminated LED light in the reward port. Only nose pokes into the reward port during this window led to reward delivery. In the Pre-training phase, there was no punishment for improper nose pokes, neither in the active port outside the tone (incorrect nose pokes) nor in the inactive port (inactive nose pokes). But in the training phase of the task, the nose pokes during the inter-trial-interval (ITI) would result in a 2-s timeout and the restart of the ITI period. For the optogenetic experiments, the photo-stimulation (1 s, 15-ms pulses, 20 Hz, 10 mW; λ = 473 nm) was delivered in the SI when the mice correctly poked the active port.

### *In vivo* fiber photometry and data analysis

To record the activities of acetylcholine *in vivo* in behaving animals, we used a commercial fiber photometry system (Inper, Hangzhou, China) to measure fluorescence signals in these neurons through an optical fiber (core diameter, 200 µm; Inper, Hangzhou, China). A patch cord (core diameter, 200 µm; Inper, Hangzhou, China) was used to connect the photometry system with the implanted optical fiber. The intensity of the blue light (λ = 470 nm) for excitation was adjusted to a low level (20∼40 µW) at the tip of the patch cord. Emitted fluorescence was bandpass filtered and focused on the sensor of a CCD camera. Photometry signals and behavioral events were aligned based on an analog TTL signal generated by the Bpod. Recorded data were exported to MATLAB for further analysis.

To correct for photobleaching of fluorescence signals (baseline drift), a bi-exponential curve was fit to the raw fluorescence trace and subtracted as follows:

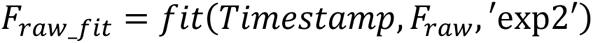

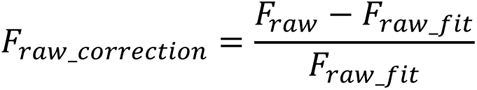

After baseline drift correction, the fluorescence signals were z-scored relative to the mean and standard deviation of signals during the baseline.

### Simultaneous fiber photometry and optogenetic stimulation

To perform simultaneous fiber photometry recording and optogenetic activation experiments, we devised a custom trigger system (based on the ThinkerTech photometry system) capable of delivering laser stimulation outside the photometry signal acquisition time window. Our photometry system employs two sampling channels: green fluorescence channel (blue LED, λ = 470 nm) and isosbestic channel (violet LED, λ = 410 nm). Laser stimulation is administered using either a blue laser (470 nm for ChR2) or a red laser (593 nm for ChrimsonR). The photometry TTL signal indicates the time window for photometry signal sampling, while the TTL trigger for laser stimulation is gated by both the control trigger from the Bpod and the low level of the photometry TTL. We sampled the fluorescence and isosbestic signals at a frequency of 20 Hz. During each 50-ms time window, we sequentially deliver blue LED (470 nm), violet LED (405 nm), and laser (blue laser, 470 nm; red laser, 593 nm) stimulation to the target area. This strategy effectively avoided any potential interference from laser artifacts on the photometry recordings.

### ROC analysis

We used the Receiver Operating Characteristic curve (ROC) to evaluate BLA ACh decodability of two trial types or outcomes. The ROC is a non-parametric statistic calculated by sweeping a classification threshold from the smallest to the largest data value and measuring the increase of single number, the area under the ROC was measured and was referred to as auROC. The statistical significance of auROC (whether it was significantly different from the chance level of 0.5) was determined using the permutation test by shuffling trial labels (1000 permutations).

### Generalized linear model (GLM)

We fit a linear encoding model for each mouse to capture the linear contribution of behavioral events to the overall observed ACh signal. The feature matrix includes representations of three behavioral events: the onset of reward, the onset of punishment, and the licking. To make the contributions of these events temporally flexible, we convolved them with a set of cubic splines. We chose splines spanning from 1 s before to 2 s after the behavior to capture both preparatory and reactive neural responses. Then we set the number of splines to seven based on three-fold cross-validation accuracy optimized on a linear encoding model. With 7 splines associated with each of the 3 behavior events, we obtained a total of 21 splines as regressors.

The encoding model takes the form y = βX + ε, where y is the photometry signal for a given mouse, with dimension T, or the number of time points. X is the design matrix composed of behavior events convolved with each cubic spline (plus one column of ones for learning the intercept term), with dimensions 22 × T. β is the set of weights, one weight for each cubic spline associated with each behavioral type plus an intercept term; the dimensions of β are 1 × 22. A percent explained by a behavioral event was expressed as the reduction of the variance in the residual responses compared to the original responses. Contributions of each behavioral event in the model were measured by the reduction of the deviance compared to a reduced model excluding the behavioral event.

## QUANTIFICATION AND STATISTICAL ANALYSIS

All statistics are indicated where used. Statistical analyses were conducted using GraphPad Prism 7 Software (GraphPad Software, CA) and MATLAB statistical toolbox (MathWorks). The statistical test used for each comparison is indicated when used. Parametric tests were used whenever possible to test differences between two or more means. Non-parametric tests were used when data distributions were non-normal. Analysis of variance (ANOVA) was used to check for main effects and interactions in experiments with repeated measures and more than one factor. All comparisons were two-tailed. Statistic hypothesis testing was conducted at a significance level of 0.05.

## SUPPLEMENTARY FIGURES, and LEGEND

**Figure S1.**
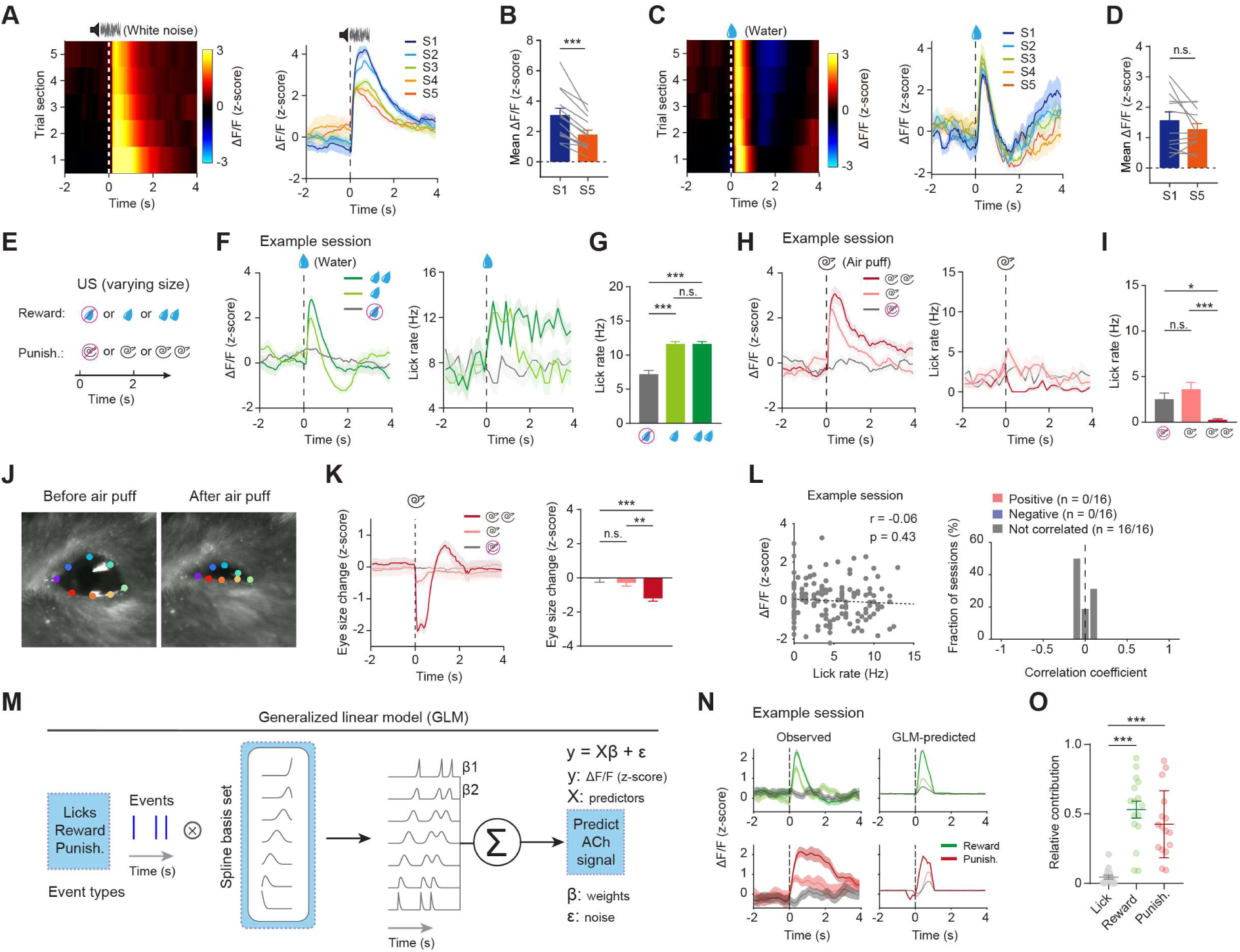
BLA ACh encodes salience but not movement. **(A)** Section-by-section (left) and average BLA ACh responses across mice in response to white noise over sections (n = 13 mice). **(B)** Quantification of BLA ACh responses following white noise delivery (0-1 s) in the first and last section (n = 13 mice; p < 0.001, paired t-test). **(C)** Section-by-section (left) and average BLA ACh responses across mice in response to water over sections (n = 12 mice). **(D)** Quantification of BLA ACh responses following water delivery (0-1 s) in the first and last section (n = 12 mice; p = 0.15, paired t-test). **(E)** Schematic of the experimental design. **(F)** Average BLA ACh responses (left) and licking rate (right) to different sizes of reward in one session. **(G)** Quantification of licking rate during reward (0-1 s) from the same session as in (F) (F_(2, 72)_ = 37.97, p < 0.001, one-way ANOVA followed by Tukey’s test). **(H)** Average BLA ACh responses (left) and licking rate (right) to different sizes of punishment in one session. **(I)** Quantification of licking rate during punishment (0-1 s) from the same session as in (h) (F_(2, 72)_ = 8.48, p < 0.001, one-way ANOVA followed by Tukey’s test). **(J)** Representative images of the eye before (left) and after (right) air puff delivery, tracked with DeepLabCut. **(K)** Average (left) and quantification (right) of the eye size change in response to different sizes of punishment (F_(2, 72)_ = 9.5, p < 0.001, one-way ANOVA followed by Tukey’s test). **(L)** Left: correlation between BLA ACh responses and licking rate in an example session (Pearson’s r = - 0.06, p = 0.43). Right: distribution of Pearson’s correlation coefficient (n = 16 sessions). **(M)** Schematic of the generalized linear model (GLM) for the relationship between behavioral events and BLA ACh responses. **(N)** Example of observed (left) and model-predicted (right) BLA ACh responses in reward (top) and punishment (bottom) trials. **(O)** Relative contribution of single model variables (lick, reward, and punishment) to model-predicted BLA ACh responses (n = 16 sessions; F_(2, 45)_ = 26.21, p < 0.001, one-way ANOVA followed by Tukey’s test). *p < 0.05, **p < 0.01, ***p < 0.001; n.s., non-significant (p > 0.05). Data are presented as mean ± SEM. Shaded areas represent SEM.

**Figure S2.**
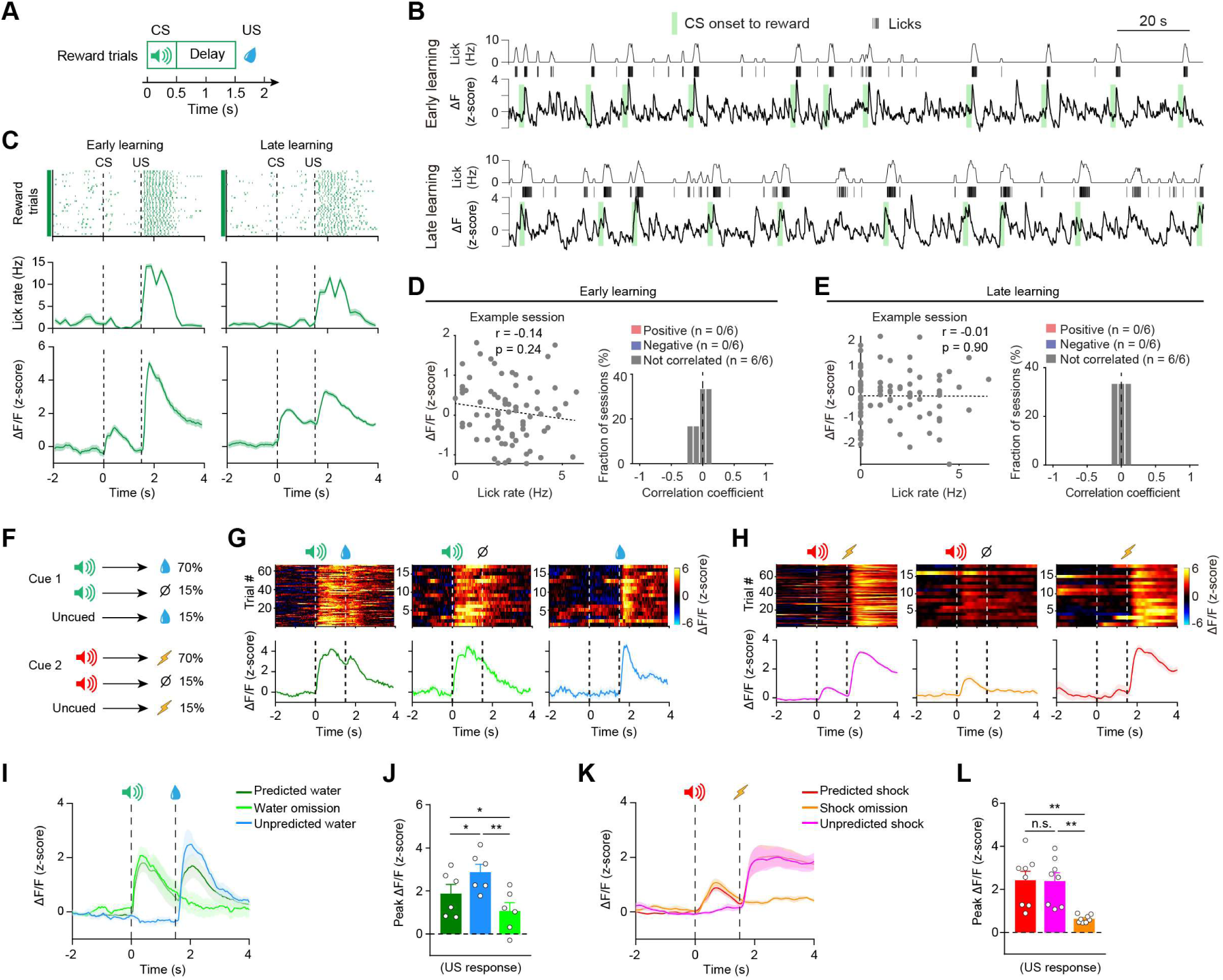
Dynamics of BLA ACh activity during learning. **(A)** Schematic of the Pavlovian conditioning task design. **(B)** Example traces of simultaneously measured behavioral and BLA ACh responses in a representative mouse at the early (top) and late (bottom) learning stages. **(C)** Top: licking events for a representative mouse at the early (left) and late (right) learning stages in the Pavlovian task. Middle: average licking rate. Bottom: average BLA ACh responses from this mouse. Dashed lines indicate the onset of CS and US. **(D, E)** Correlation between licking rate and BLA ACh responses during baseline in an example session (left) and the distribution of Pearson’s correlation coefficient (right) at the early (D) (Pearson’s r = −0.14, p = 0.24) and the late (E) (Pearson’s r = −0.01, p = 0.90) learning stages (n = 6 mice). **(F)** Schematics for the probabilistic Pavlovian conditioning task with cue and outcome probabilities indicated. **(G)** Trial-by-trial (top) and average (bottom) responses of BLA ACh in different types of reward trials from an example session. **(H)** Trial-by-trial (top) and average (bottom) responses of BLA ACh in different types of punishment trials from an example session. **(I)** Average BLA ACh responses (n = 6 mice) in different types of reward trials. **(J)** Quantification of the peak BLA ACh responses in different types of reward trials (F_(2, 15)_ = 5.19, p = 0.019, one-way ANOVA followed by Tukey’s test). **(K)** Average BLA ACh responses (n = 8 mice) in different types of punishment trials. **(L)** Quantification of the peak BLA ACh responses in different types of punishment trials (F_(2, 21)_ = 9.73, p = 0.001, one-way ANOVA followed by Tukey’s test). *p < 0.05, **p < 0.01; n.s., non-significant (p > 0.05). Data are presented as mean ± SEM. Shaded areas represent SEM.

**Figure S3.**
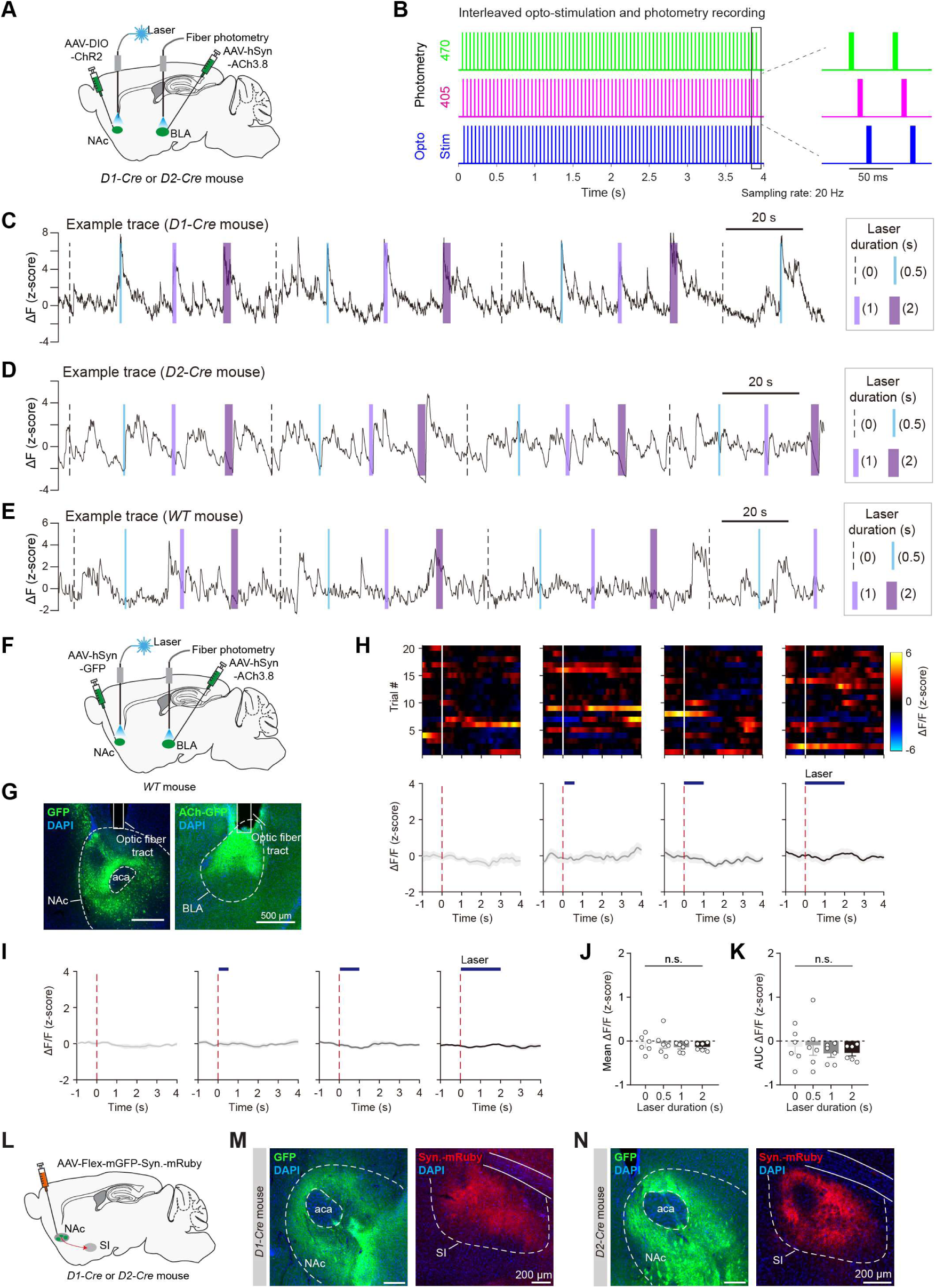
Laser stimulation of the NAc does not affect BLA ACh in GFP mice. **(A)** Schematic of the approach to activate the NAc and record BLA ACh simultaneously. **(B)** Diagram showing simultaneous opto-stimulation and photometry recording interleaved at the frequency of 20 Hz. **(C-E)** Example trace of BLA ACh responses with laser stimulation from a representative *D1-Cre* mouse (C), *D2-Cre* mouse (D), and wild-type (WT) mouse (E). **(F)** Schematic of the strategy to record BLA ACh while delivering laser into the NAc of a WT mouse. **(G)** Left: representative image of GFP expression in the NAc. Right: representative image of ACh3.8 expression in the BLA. **(H)** Trial-by-trial heatmap (top) and average (bottom) responses of BLA ACh while delivering laser into the NAc with different durations in an example session of a WT mouse. **(I)** Average BLA ACh responses (n = 6) while delivering the laser into the NAc with different laser durations in WT mice. **(J, K)** Quantification of the mean (J) (F_(3, 20)_ = 0.36, p = 0.78) and AUC (K) (F_(3, 20)_ = 0.38, p = 0.77) of BLA ACh signals while delivering laser into the NAc with different laser durations in WT mice (n = 6). One-way ANOVA. **(L)** Schematic of anterograde tracing of NAc D1 or D2 MSNs. **(M, N)** Example confocal images of the injection site (soma; green) in the NAc (left) and the axonal terminals (red) in the SI (right) from a representative *D1-Cre* (M) or *D2-Cre* (N) mouse. n.s., non-significant (p > 0.05). Data are presented as mean ± SEM. Shaded areas represent SEM.

**Figure S4.**
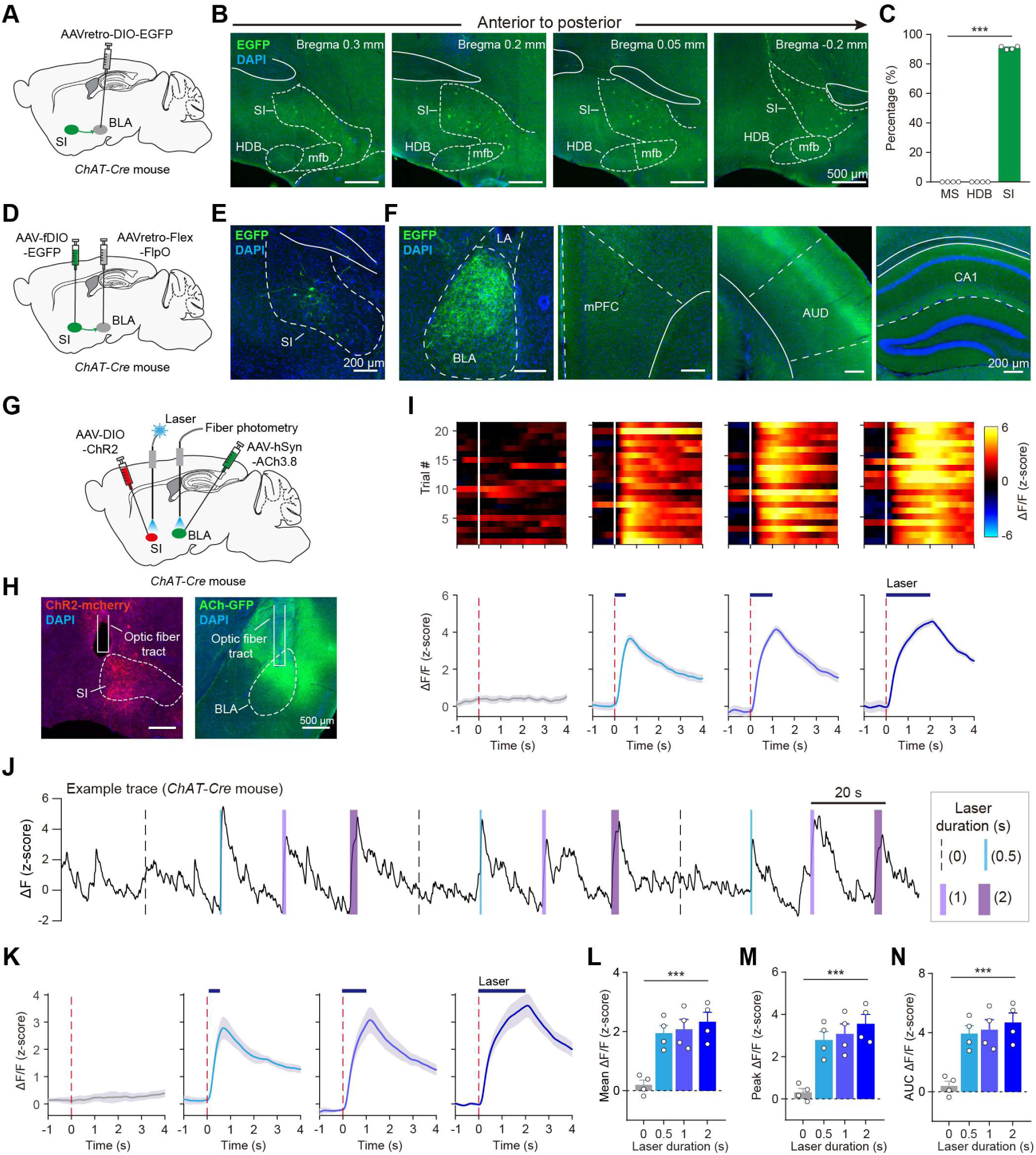
Activation of SI ChAT neurons induces potent ACh release in the BLA. **(A)** Schematic of retrograde labeling of ChAT neurons that project to the BLA. **(B)** Representative images showing labeled ChAT neurons in the basal forebrain. **(C)** Quantification of labeled ChAT neurons in the medial septum (MS), diagonal band of Broca (HDB), and SI (n = 4; F_(2, 9)_ = 25408, p < 0.001). One-way ANOVA followed by Tukey’s test. **(D)** Schematic of anterograde tracing of BLA-projecting SI ChAT neurons. **(E, F)** Representative images showing EGFP-expressing cell bodies in SI (E) and their labeled axonal downstream (F). **(G)** Schematic of the strategy to record BLA ACh while activating SI ChAT neurons. **(H)** Left: representative image of ChR2 expression in SI ChAT neurons. Right: representative image of Ach3.8 expression in the BLA. **(I)** Trial-by-trial (top) and average (bottom) responses of BLA ACh while activating SI ChAT neurons with different laser durations in an example session. **(J)** Example trace of BLA ACh responses when activating SI ChAT neurons with different laser durations. **(K)** Average BLA ACh responses (n = 4 mice) while delivering the laser into the SI with different laser durations. **(L-N)** Quantification of the mean (L) (F_(3, 12)_ = 12.22, p < 0.001), peak (M) (F_(3, 12)_ = 13.59, p < 0.001), and AUC (N) (F_(3, 12)_ = 12.22, p < 0.001) of BLA ACh signals while activating SI ChAT neurons with different laser durations (n = 4 mice). One-way ANOVA. ***p < 0.001. Data are presented as mean ± SEM. Shaded areas represent SEM.

**Figure S5.**
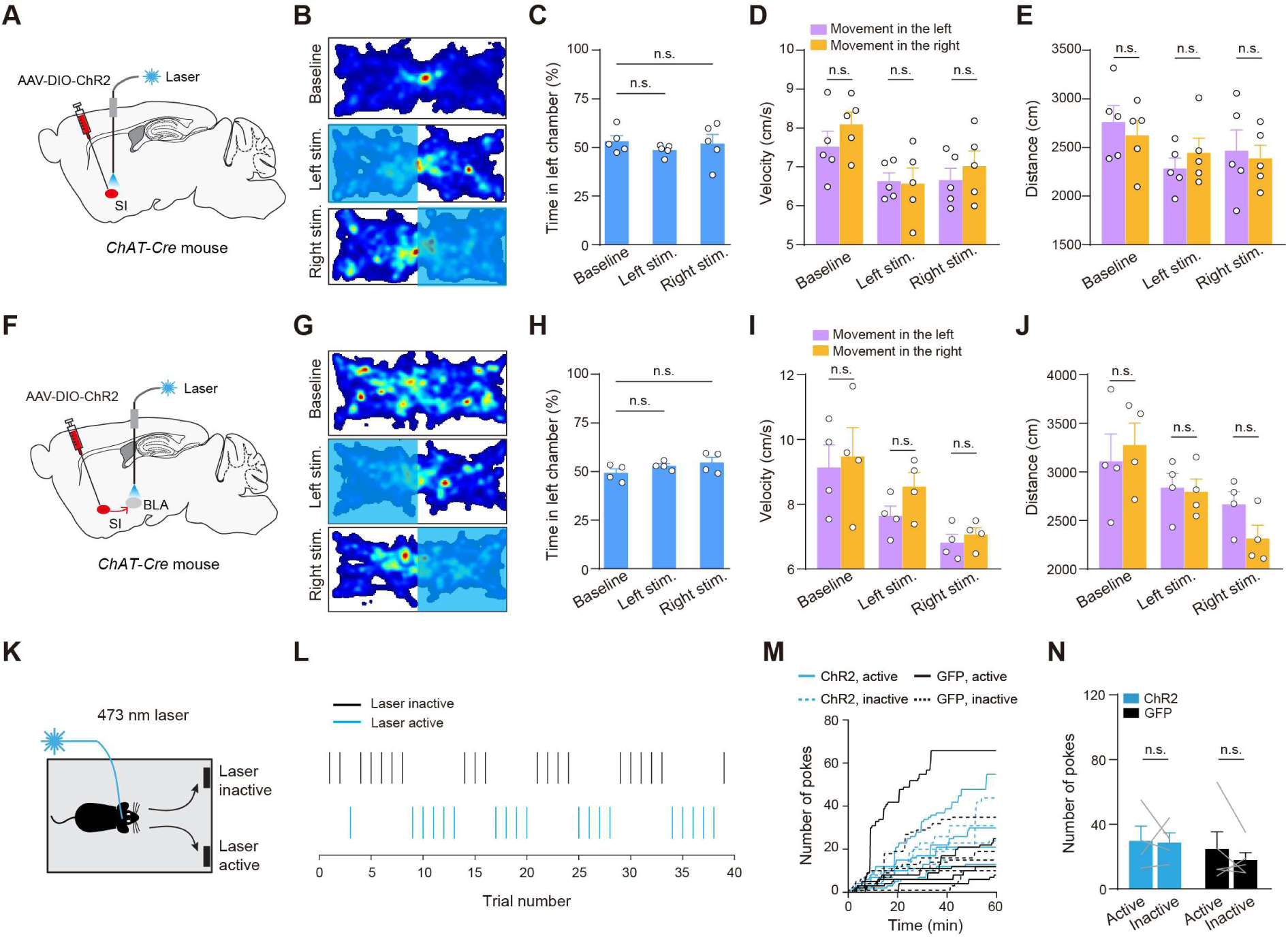
Activation of SI^ChAT^ neurons and SI^ChAT^➔BLA pathway does not induce valence or drive reinforcement. **(A)** Schematic of the approach for optogenetic activation of SI^ChAT^ neurons. **(B)** Heatmaps for the activity of a representative mouse at baseline (top), or in a situation where entering the left (middle) or right (bottom) side of the chamber triggered photo-stimulation in the SI. **(C-E)** Quantification of the mouse activity as shown in (B), showing that activation of SI^ChAT^ neurons does not induce place preference or aversion (C) (F_(2, 12)_ = 0.54, p = 0.59, one-way ANOVA followed by Tukey’s test), affect movement velocity (D) (F_(2, 12)_ = 0.74, p = 0.50, two-way ANOVA followed by Bonferroni’s test), or affect distance traveled (E) (F_(2, 12)_ = 0.44, p = 0.66, two-way ANOVA followed by Bonferroni’s test). **(F)** Schematic of the approach for optogenetic activation of SI^ChAT^➔BLA pathway. **(G)** Heatmaps for the activity of a representative mouse at baseline (top), or in a situation where entering the left (middle) or right (bottom) side of the chamber triggered photo-stimulation of SI^ChAT^➔BLA pathway. **(H-J)** Quantification of the mouse activity as shown in (G), showing that activation of SI^ChAT^➔BLA pathway does not induce place preference or aversion (H) (F_(2, 9)_ = 1.73, p = 0.23, one-way ANOVA followed by Tukey’s test), affect movement velocity (I) (F_(2, 9)_ = 1.30, p = 0.32, two-way ANOVA followed by Bonferroni’s test), or affect distance traveled (J) (F_(2, 9)_ = 1.77, p = 0.23, two-way ANOVA followed by Bonferroni’s test). **(K)** Schematic of the self-stimulation test. **(L-N)** Optogenetic activation of SI^ChAT^➔BLA does not support self-stimulation. **(L)** Poking events at one port where poking triggered the photoactivation of SI^ChAT^➔BLA (active), and the other port where poking did not trigger the photo-stimulation (inactive) from an example session of a ChR2 mouse. **(M)** Cumulative curves for the poking responses at a port where poking triggered the photo-stimulation (active), and a port where poking did not trigger the photo-stimulation (inactive), in ChR2 mice (n = 4), or WT mice (as the control; n = 5). **(N)** Quantification of the poking responses as shown in (M) (F_(1, 7)_ = 0.22, p = 0.66, two-way ANOVA followed by Bonferroni’s test). n.s., non-significant (p > 0.05). Data are presented as mean ± SEM.

**Figure S6.**
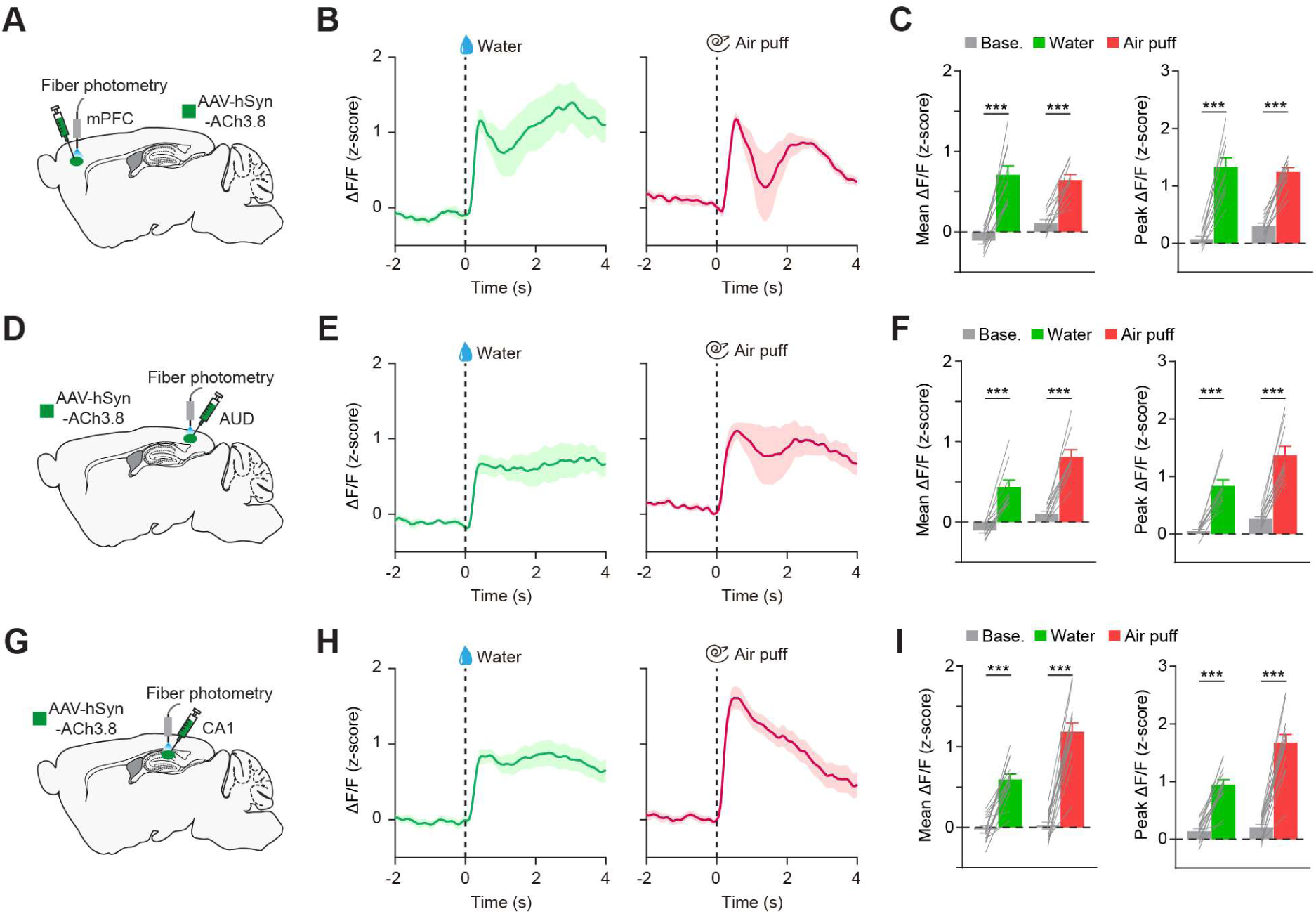
The responses of cortical and hippocampal ACh to reward and punishment. **(A)** Schematic of ACh recording in the mPFC. **(B)** Average response of mPFC ACh to water (left) and air puff (right) from a representative mouse. **(C)** Quantification of the mean (left) and peak (right) mPFC ACh responses to water and air puff (p < 0.001, paired t-test). **(D)** Schematic of ACh recording in the AUD. **(E)** Average response of AUD ACh to water (left) and air puff (right) from a representative mouse. **(F)** Quantification of the mean (left) and peak (right) AUD ACh responses to water and air puff (p < 0.001, paired t-test). **(G)** Schematic of ACh recording in the CA1. **(H)** Average response of CA1 ACh to water (left) and air puff (right) from a representative mouse. **(I)** Quantification of the mean (left) and peak (right) CA1 ACh responses to water and air puff (p < 0.001, paired t-test). ***p < 0.001. Data are presented as mean ± SEM. Shaded areas represent SEM.

**Figure S7.**
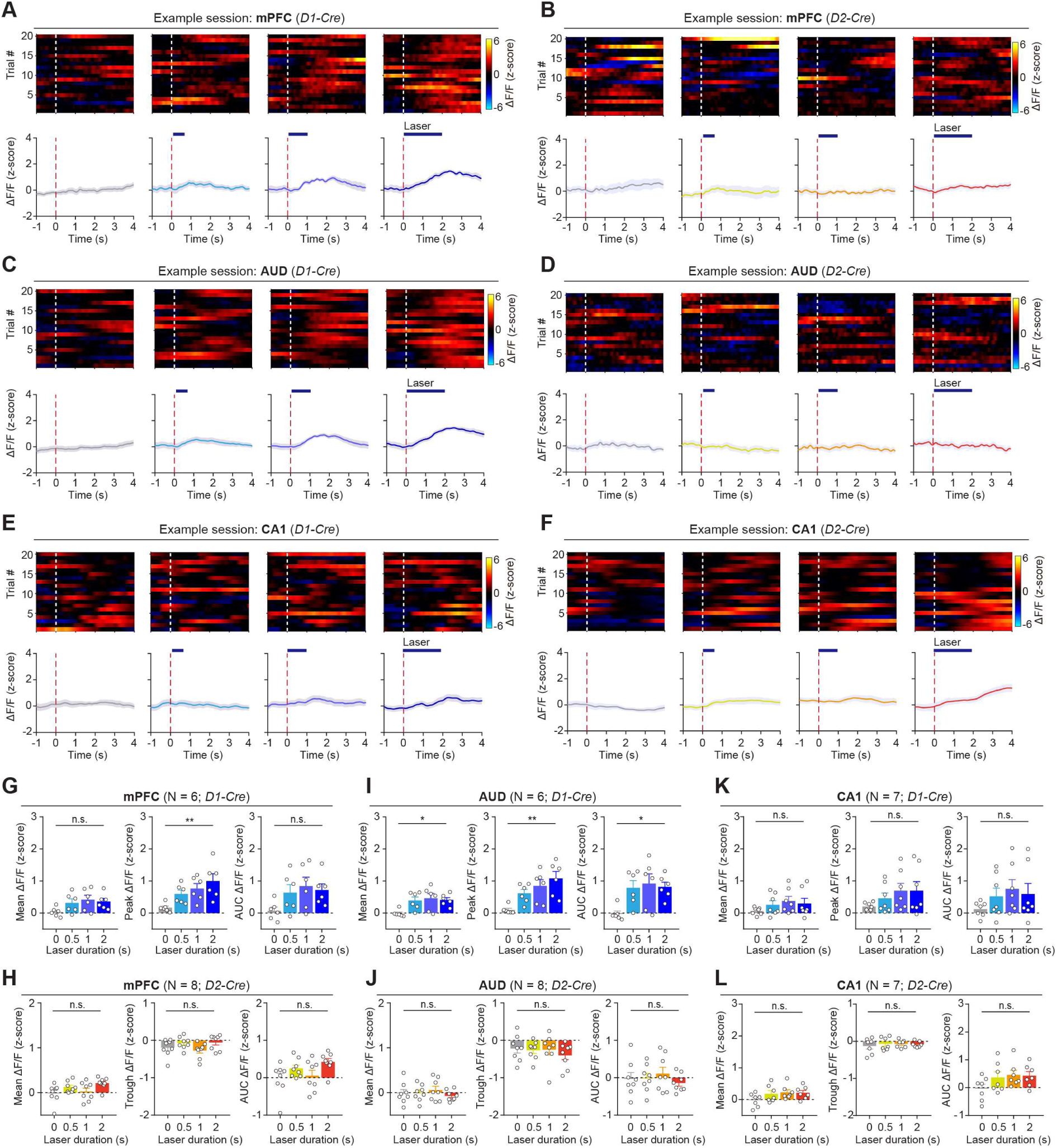
Activation of NAc➔SI pathway does not affect ACh release in the cortex and hippocampus. **(A, B)** Trial-by-trial heatmap (top) and average (bottom) ACh responses in the mPFC while activating NAc^D1^➔SI (A) and NAc^D2^➔SI (B) with different laser durations in an example session. **(C, D)** Trial-by-trial heatmap (top) and average (bottom) ACh responses in the AUD while activating NAc^D1^➔SI (C) and NAc^D2^➔SI (D) with different laser durations in an example session. **(E, F)** Trial-by-trial heatmap (top) and average (bottom) ACh responses in the CA1 while activating NAc^D1^➔SI (E) and NAc^D2^➔SI (F) with different laser durations in an example session. **(G, H)** Quantification of the mean (left), peak (middle) and AUC (right) of mPFC ACh responses while activating NAc^D1^➔SI (G) (left, F_(3, 20)_ = 2.41, p = 0.097; middle, F_(3, 20)_ = 5.34; p = 0.0072; right, F_(3, 20)_ = 2.43, p = 0.095) and NAc^D2^➔SI (H) (left, F_(3, 28)_ = 2.81, p = 0.058; middle, F_(3, 28)_ = 2.14, p = 0.12; right, F_(3, 28)_ = 2.81, p = 0.058) with different laser durations. One-way ANOVA. **(I, J)** Quantification of the mean (left), peak (middle) and AUC (right) of AUD ACh responses while activating NAc^D1^➔SI (I) (left, F_(3, 20)_ = 4.50, p = 0.014; middle, F_(3, 20)_ = 6.85, p = 0.0023; right, F_(3, 20)_ = 4.49, p = 0.015) and NAc^D2^➔SI (J) (left, F_(3, 28)_ = 0.54, p = 0.66; middle, F_(3, 28)_ = 0.44, p = 0.72; right, F_(3, 28)_ = 0.52, p = 0.67) with different laser durations. One-way ANOVA. **(K, L)** Quantification of the mean (left), peak (middle) and AUC (right) of CA1 ACh responses while activating NAc^D1^➔SI (K) (left, F_(3, 24)_ = 1.06, p = 0.38; middle, F_(3, 24)_ = 1.35, p = 0.28; right, F_(3, 24)_ = 1.06, p = 0.38) and NAc^D2^➔SI (L) (left, F_(3, 24)_ = 1.78, p = 0.18; middle, F_(3, 24)_ = 0.24, p = 0.87; right, F_(3, 24)_ = 1.81, p = 0.17) with different laser durations. One-way ANOVA. *p < 0.05, **p < 0.01; n.s., non-significant (p > 0.05). Data are presented as mean ± SEM. Shaded areas represent SEM.

**Figure S8.**
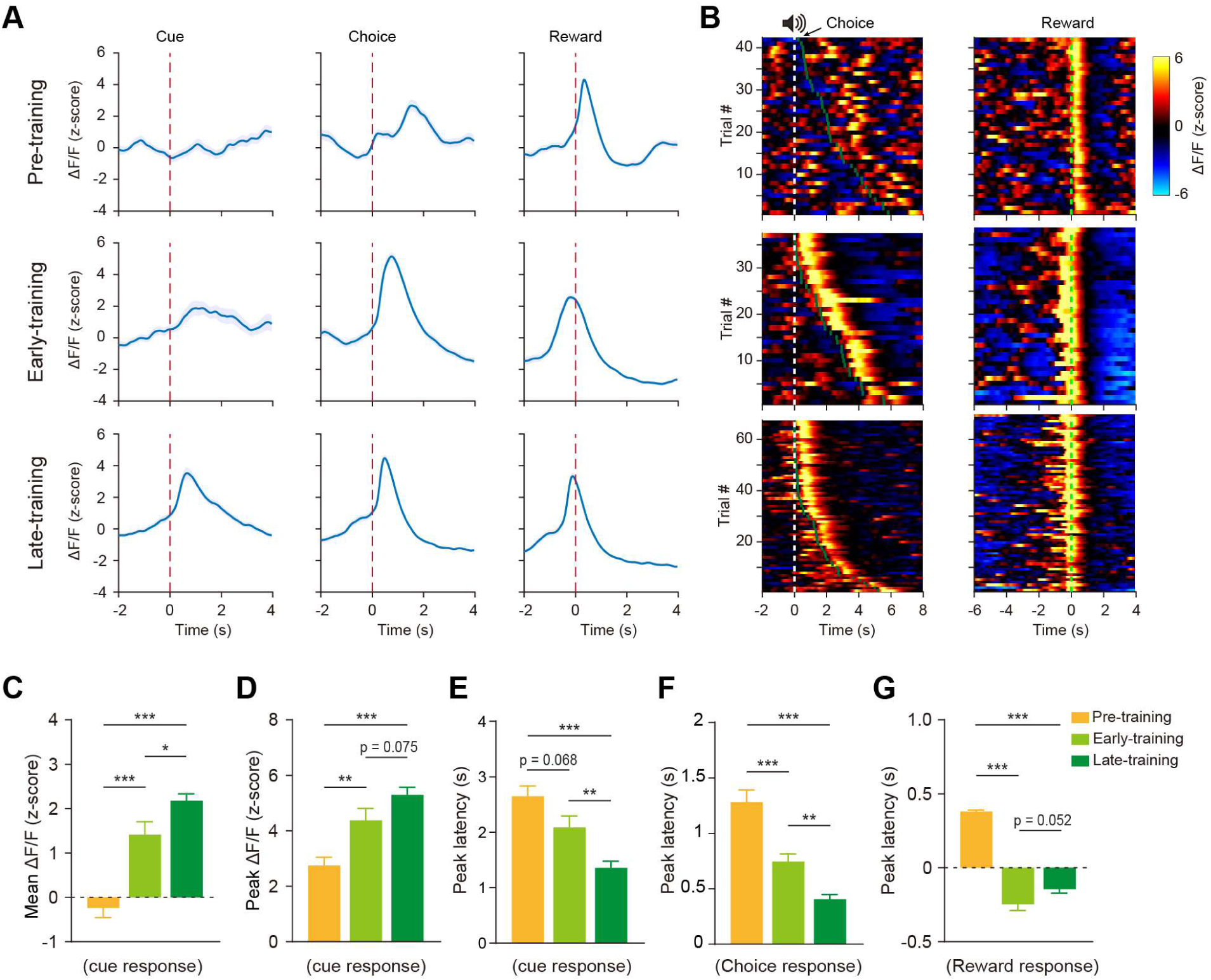
BLA ACh responses shift from US to CS during cue-reward learning. **(A)** Average BLA ACh responses aligned to the cue onset (left), initial poke (middle), and reward retrieval (right) in the pre-training (top), early-training (middle), and late-training (bottom) stages of the cue-reward learning task from a representative mouse. **(B)** Trial-by-trial heatmap of BLA ACh responses aligned to the cue onset (left) and reward retrieval (right) at the pre-training (top), early-training (middle), and late-training (bottom) stages from the same mouse as in (A). White dashed line, the cue onset; dark green dashed line, the initial poke; green dashed line, the reward retrieval. **(C, D)** Quantification of the mean (C) (F_(2, 143)_ = 36.74, p < 0.001) and peak (D) (F_(2, 143)_ = 16.24, p < 0.001) of BLA ACh responses to the cue at different training stages. One-way ANOVA followed by Tukey’s test. **(E-G)** Quantification of the peak latency of BLA ACh responses to the cue (E) (F_(2, 143)_ = 17.98, p < 0.001), initial poke (F) (F_(2, 143)_ = 39.83, p < 0.001), and reward retrieval (G) (F_(2, 143)_ = 107.2, p < 0.001) at different training stages. One-way ANOVA followed by Tukey’s test. *p < 0.05, **p < 0.01, ***p < 0.001. Data are presented as mean ± SEM. Shaded areas represent SEM.

**Figure S9.**
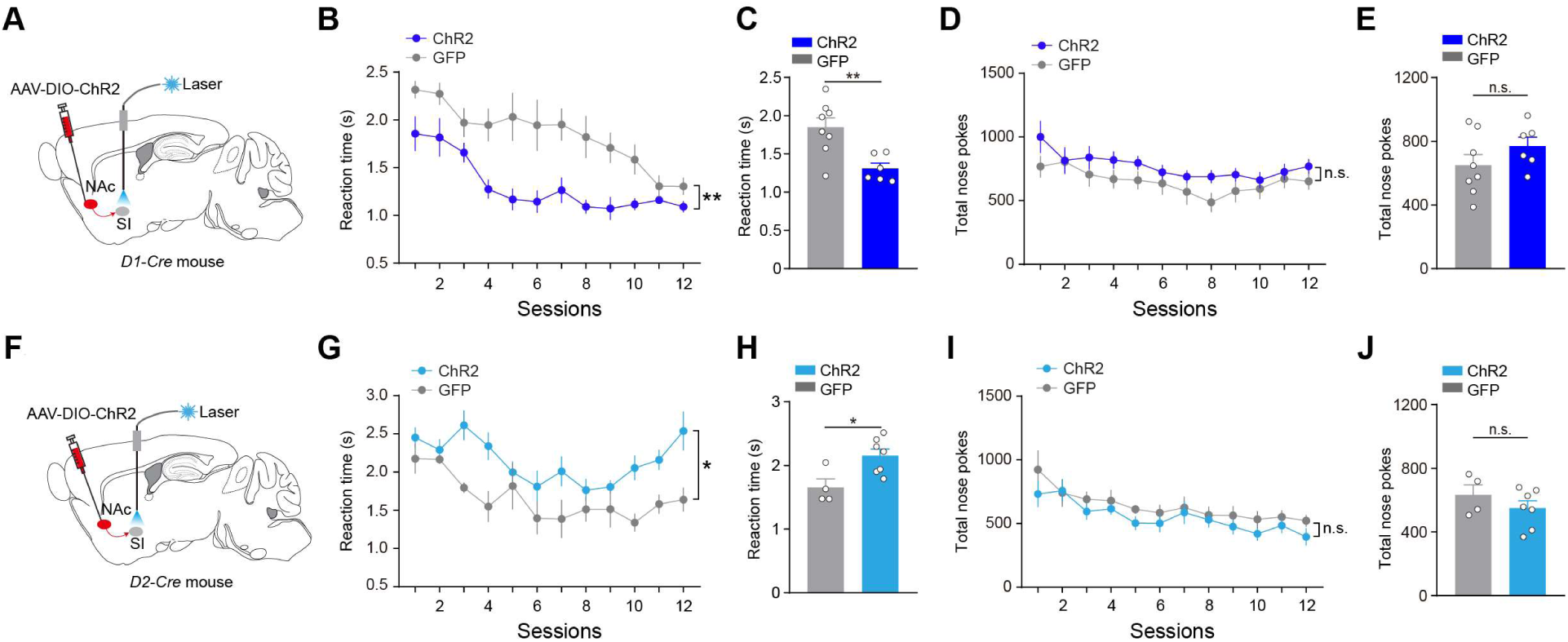
Activation of NAc^D1^➔SI and NAc^D2^➔SI pathways affect reaction time but do not impair movement. **(A)** Schematic of the approach to activate NAc^D1^➔SI pathway. **(B, C)** Reaction time of the ChR2 (n = 6) and GFP (n = 8) mice in each session (B) (interaction, F_(11, 132)_ = 1.90, p = 0.044; group, F_(1, 12)_ = 11.78, p = 0.0050; two-way ANOVA), and average across sessions (C) (p = 0.0050, t-test) with closed-loop stimulation of NAc^D1^➔SI in the cue-reward learning task. **(D, E)** Number of total nose pokes of the ChR2 (n = 6) and GFP (n = 8) mice in each session (D) (interaction, F_(11, 132)_ = 0.69, p = 0.75; group, F_(1, 12)_ = 1.70, p = 0.22; two-way ANOVA), and average across sessions (E) (p = 0.22, t-test) with closed-loop stimulation of NAc^D1^➔SI in the cue-reward learning task. **(F)** Schematic of the approach to activate NAc^D2^➔SI pathway. **(G, H)** Reaction time of the ChR2 (n = 7) and GFP (n = 4) mice in each session (G) (interaction, F_(11, 99)_ = 1.62, p = 0.104; group, F_(1, 9)_ = 8.36, p = 0.018; two-way ANOVA), and average across sessions (H) (p = 0. 018, t-test) with closed-loop stimulation of NAc^D2^➔SI in the cue-reward learning task. **(I, J)** Number of total nose pokes of the ChR2 (n = 7) and GFP (n = 4) mice in each session (I) (interaction, F_(11, 99)_ = 0.40, p = 0.95; group, F_(1, 9)_ = 1.15, p = 0.31; two-way ANOVA), and average across sessions (J) (p = 0.31, t-test) with closed-loop stimulation of NAc^D2^➔SI in the cue-reward learning task. *p < 0.05, **p < 0.01; n.s., non-significant (p > 0.05). Data are presented as mean ± SEM.

